# Population encoding of stimulus features along the visual hierarchy

**DOI:** 10.1101/2023.06.27.545450

**Authors:** Luciano Dyballa, Andra M. Rudzite, Mahmood S. Hoseini, Mishek Thapa, Michael P. Stryker, Greg D. Field, Steven W. Zucker

**Author notes:** Corresponding author Steven W. Zucker,.

## Abstract

The retina and primary visual cortex (V1) both exhibit diverse neural populations sensitive to diverse visual features. Yet it remains unclear how neural populations in each area partition stimulus space to span these features. One possibility is that neural populations are organized into discrete groups of neurons, with each group signaling a particular constellation of features. Alternatively, neurons could be continuously distributed across feature-encoding space. To distinguish these possibilities, we presented a battery of visual stimuli to mouse retina and V1 while measuring neural responses with multi-electrode arrays. Using machine learning approaches, we developed a manifold embedding technique that captures how neural populations partition feature space and how visual responses correlate with physiological and anatomical properties of individual neurons. We show that retinal populations discretely encode features, while V1 populations provide a more continuous representation. Applying the same analysis approach to convolutional neural networks that model visual processing, we demonstrate that they partition features much more similarly to the retina, indicating they are more like big retinas than little brains.

---

The output of the mouse retina is formed by a set of about 40 distinct types of retinal ganglion cells (RGCs) [7, 41]. These RGC types exhibit distinct morphologies, gene expression profiles, and visual responses. This has generated a coherent perspective on retinal output: distinct cell types signal distinct visual features to downstream brain areas. How are these distinct features organized in the cortex? Many studies have focused on parallel pathways in sensory systems and in the visual system in particular [39, 43]. One possibility is that the visual features signaled by parallel pathways originating from the different RGC classes would be combined to produce new groupings of features in the cortex [59, 12]. An alternative possibility is that visual cortical organization is a continuum, fundamentally different from that of the retina, despite the mounting evidence for distinct neuronal identities in V1, with cortical cell types of different morphologies, transcriptomes, and intrinsic physiological properties [102].

To determine whether stimulus features are represented similarly over the parallel pathways between retina and visual cortex, we have devised the “neural encoding manifold”—a novel analysis of the organization of neurons signaling a wide range of visual response features. We have applied it to both retina and V1 using, for the first time, matched stimulus ensembles across the two populations of neurons. Our conclusion is that the population-level representations of stimulus features are fundamentally different between retina and visual cortex, the former organized as distinct clusters while the latter as a continuum.

## An ensemble of grating and flow stimuli

To identify the degree of similarity between population-level neural representations of stimuli in the retina and V1, we performed multi-electrode array (MEA) recordings in both areas utilizing matched stimuli and analysis procedures. MEAs were used to measure the responses of mouse RGCs *ex vivo*, and responses of mouse V1 neurons *in vivo* (Fig. 1a,b). Sinusoidal drifting gratings and optical flow stimuli were presented at matched spatial frequencies and contrasts in the retina and V1 experiments (Fig. 1c). Drifting gratings were presented at two spatial frequencies and drifted in eight directions. Flow stimuli were presented at two contrasts (positive and negative), consisting of dots and oriented line segments, and drifted in eight directions. Flow stimuli were chosen because they mimic certain features of naturalistic stimuli, and previous work has shown that they engage nonlinearities in V1 that are not predicted based on the responses to gratings [35]. In addition, flow and grating stimuli were spatially isotropic (or repeating), allowing us to study response properties independent of receptive field location. Individual neuronal responses were summarized by a two-dimensional response map, where each row shows the response to a particular stimulus direction of motion; one such map was produced for each neuron and for each of the six stimuli presented (Fig. 1d).

**Fig. 1:**
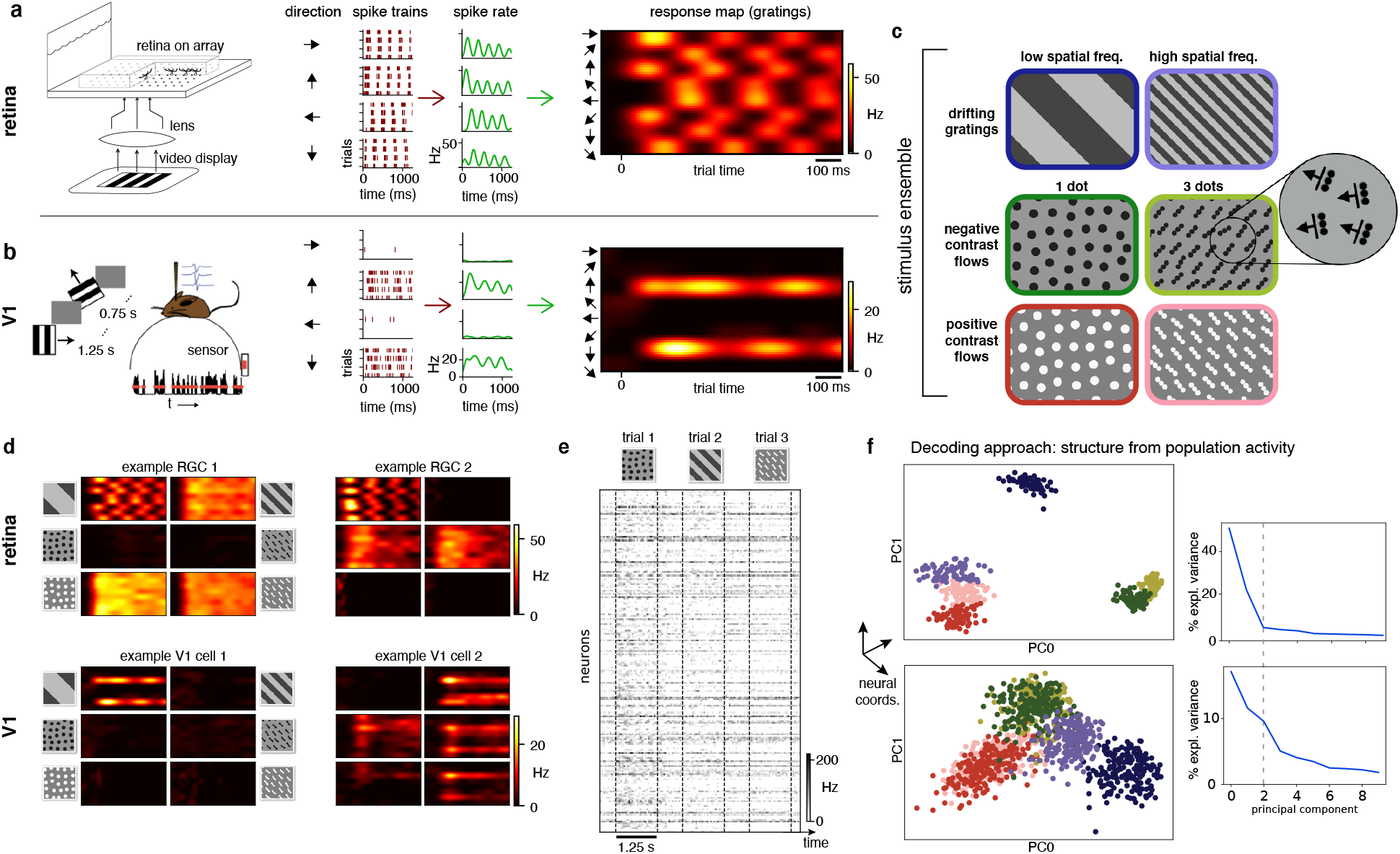
Experimental set-up and stimulus ensemble. **a–b**, MEA measurements from mouse RGCs and V1 neurons. Spikes from drifting stimuli are trial-averaged and collected into response maps. Examples shown from an ON-center RGC (**a**) and an orientation-selective cell in V1 (**b**). **c**, The stimulus ensemble consists of low and high spatial frequency gratings and positive or negative contrast flow patterns composed of either single dots or 3-dot oriented line-segments moving in 8 directions separated by 45*^◦^*(see Methods). The illustrated 3 *×* 2 arrangement will be used throughout. **d**, Two example RGC (top) and V1 (bottom) response profiles illustrating the diverse response dynamics to the different stimuli. Left cells identical to (**a, b**). **e**, Rasters showing population responses: neurons *×* spike trains. **f**, *Left:* Example of standard embedding approaches [84, 32, 80, 92] arranges trials in neural coordinates and enables “decoding”, i.e., inferring the stimulus from the neural state (top is retina, bottom is V1). Each point represents the response of the neural population to a given stimulus on a trial in a low-dimensional space determined by principal components analysis (see Methods); colors indicate the different stimuli. *Right:* Principal value spectrum associated with the principal components for RGCs (top) and V1 (bottom): although stimulus decoding is clear in both, cortex appears to require more dimensions to capture variability in the responses.

## Organizing neurons, stimuli, and responses via the encoding manifold

To investigate and compare the organization of retinal and V1 population responses, we sought to organize the population responses on a manifold. Previous work has taken a “stimulus perspective” on these responses [32, 27, 60, 80, 22, 67], producing a low-dimensional representation that organizes stimuli by the population response to each trial (Fig. 1e), allowing one to decode the identity of the stimulus that gave rise to the constellation of responses to a single stimulus trial. While these approaches are useful, our goal was distinct; we sought to identify how individual neurons contributed to signaling different stimulus features with respect to the entire neural population. Thus, we switched from the stimulus-based “decoding” perspective used previously (Fig. 1e,f) to a neuron-based “encoding” perspective by developing a novel “encoding manifold” (Fig. 2). Each point on this encoding manifold is a neuron, not a stimulus, so it reveals how neurons are distributed in stimulus/response space. Thus neurons are organized by how they respond to features within the stimulus ensemble. These features included positive versus negative stimulus contrasts, high versus low spatial frequencies, motion in different directions, orientation, and any other features present in the stimulus set that drive different responses across the neural population. Neurons with similar feature selectivity and response dynamics will be nearby on the manifold, while neurons responding to different stimulus features or with distinct dynamics will be far apart on the manifold. Thus, the axes of this space are related to both stimulus features and temporal response characteristics. This approach differs substantially from the conventional one, which focuses solely on the stimulus selectivity of neurons. Here, we give equal emphasis on the dynamics of visual responses as to their stimulus selectivity. This dual focus is important for understanding how the brain processes visual input because the differences in dynamics are sufficiently large (tens of ms) that processing in higher visual areas will be strongly affected [75].

**Fig. 2:**
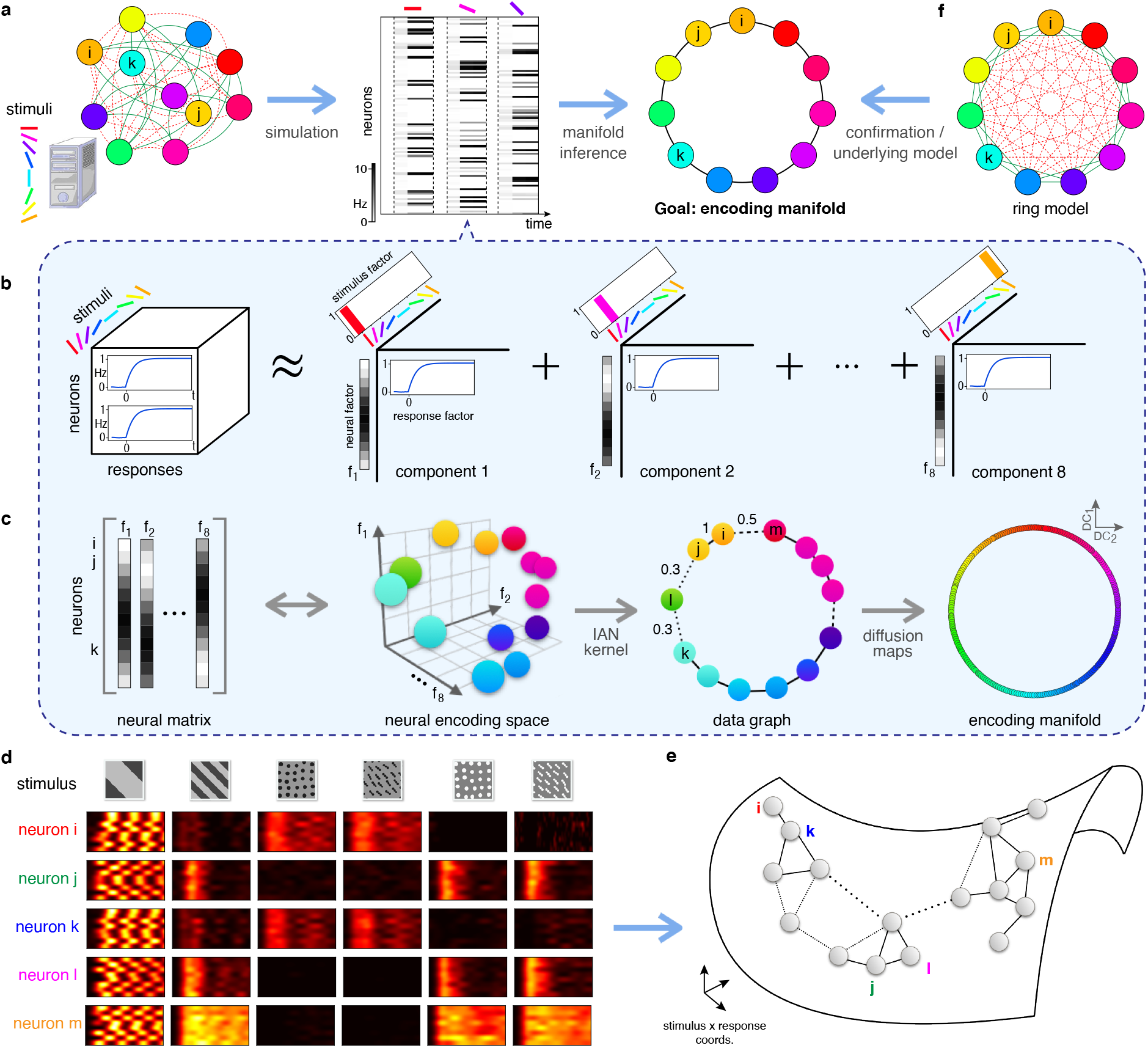
Illustration of the encoding manifold algorithm applied to an artificial example. **a**, Suppose we are given an unknown circuit of orientation-selective neurons *(left)*. Stimulating it with orientated gratings yields responses; a spike train for each neuron for each stimulus trial *(middle)*. The goal is to use these data in an unsupervised manner to organize the neurons into a manifold: each point on the manifold is a neuron and nearby neurons respond similarly in time to the stimulus ensemble *(right)*. **b–c**, The manifold inference proceeds in two stages. **b**, Stage 1: non-negative tensor factorization [97] is used to multi-cluster groups of neurons by stimuli and by response. For this artificial example all temporal responses are identical, so different neurons are organized by the different stimuli. Since 8 orientations are used, 8 latent components emerge (a procedure for selecting the number of components is described in Methods). **c**, Stage 2: The neural factors are gathered into a matrix whose dimensions are #neurons *×* #factors; a row in this matrix represents the loading of each neuron in each factor *f_i_, i* = 1, 2*, …,* 8. Equivalently, each row defines a vector in a stimulus/response-space, encoding the response of each neuron to each stimulus. A weighted data graph is then built from an iterated adaptive neighborhood similarity kernel [34] in this space and used with diffusion maps for manifold inference [26, 25]. For this artificial example the manifold is a ring in which neurons tuned to nearby orientations are neighbors. **f**, This provides insight into the artificial problem we set up, namely the artificial ring model of orientation tuning from [13]. **d–e**, Actual data are more complicated. To illustrate conceptually, five example RGCs and their responses to the stimuli are shown (**d**). Note *i* and *k* respond similarly, so they should be close on the encoding manifold (**e**); neurons *i* and *j* respond differently, so they are distant on the manifold. Although *m* responds to the same stimuli as *j* and *l*, its response dynamics are different, so it is distant from the others. The underlying data graph is shown superimposed on the manifold.

To illustrate the encoding manifold approach and validate that it (1) recovers the underlying functional structure of a neural population and (2) organizes these neurons appropriately on a manifold, we used the popular artificial ring model of orientation tuning [13, 78] (Fig. 2f; see Methods). In this model, neurons tuned to nearby orientations excite one another while those tuned to different orientations inhibit each other. Equilibria of the network are the points on a ring (the space of possible orientations). We simulated responses of artificial neurons to low frequency gratings drifting in eight different directions (Fig. 2a), each with the same dynamics. The encoding manifold was constructed in a two-stage procedure with no knowledge of the model’s underlying circuitry (Fig. 2b,c; see Methods): first, non-negative tensor factorization [97] was used to relate stimuli to neural responses across the population of recorded cells. A neural encoding space was then constructed by bundling the neural factors into a matrix, in which a similarity kernel [34] could be defined. From this, diffusion maps—a non-linear inference algorithm [26, 25]—yielded the manifold. Expressed in diffusion coordinates, the ring emerged solely from the responses, even though the algorithm knew nothing of the underlying circuit organization.

In general, neural data will be more complex than that produced by a simple ring model (Fig. 2d), with many neurons exhibiting mixed selectivity [77, 35]. As such, the manifold is likely to be richer than the ring, but the idea generalizes: neurons with similar response profiles will be nearby on the manifold. A population of neurons with distinct responses will be far from other neurons on the manifold (Fig. 2e).

## Comparing retina and V1

We first applied the encoding manifold approach to MEA recordings from 1149 mouse RGCs (Fig. 3). Results from individual retinas were sufficiently similar that data from three retinas were combined to embed all data at once (see Fig. S3d). The resulting manifold exhibited clusters of neurons, with the cells in each cluster exhibiting similar response dynamics to similar stimuli. Other clusters, with different stimulus/response profiles, were relatively separated, so that following a trajectory along the manifold would reveal a population of nearly similar cells followed by an abrupt transition to another, different group of cells. See Discussion for further analysis of this manifold.

**Fig. 3:**
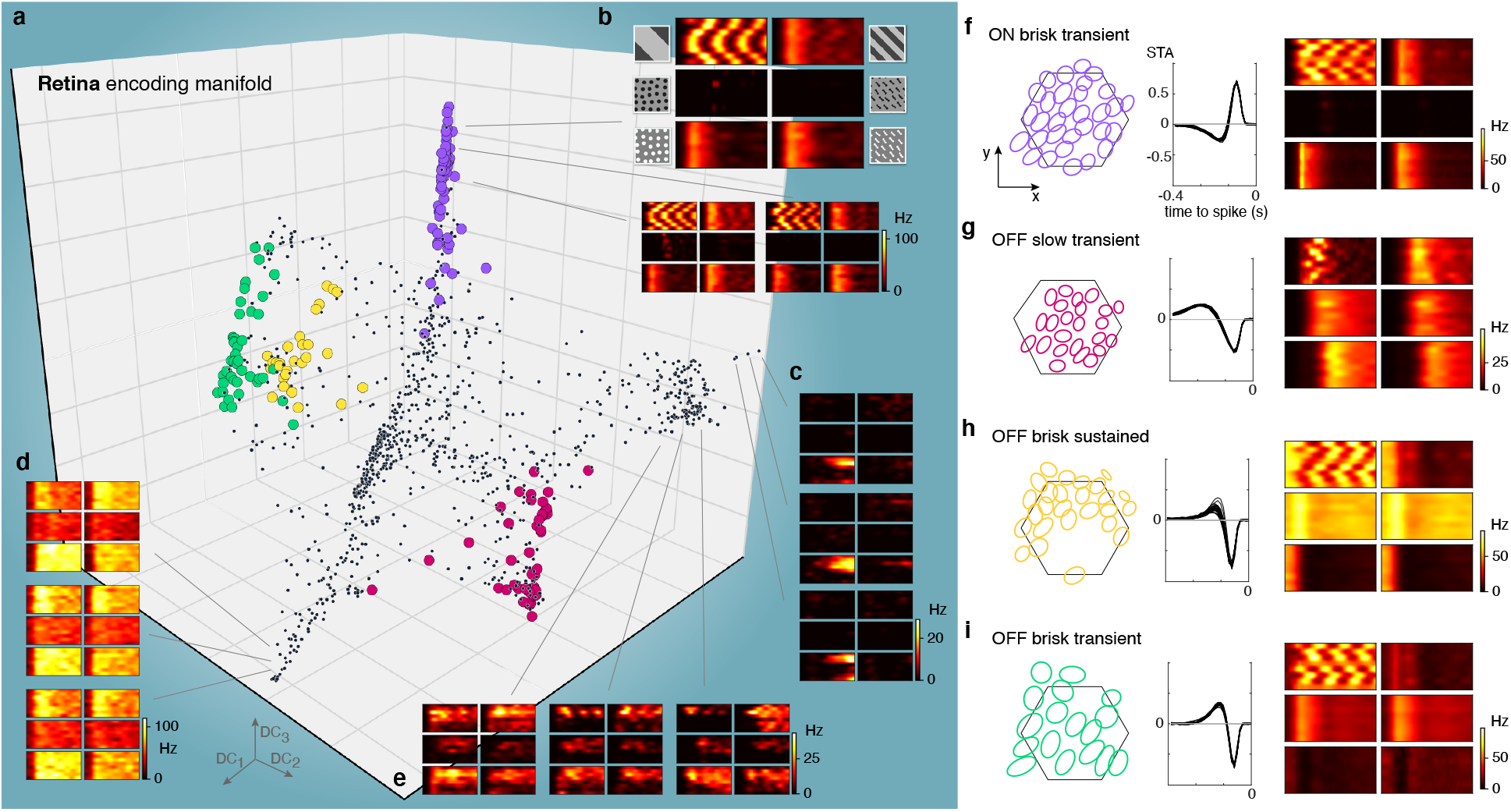
The encoding manifold formed by RGCs is clustered and organized by cell types. **a**, Projection of the encoding manifold formed by RGCs onto the first three diffusion coordinates (*n* = 1149 RGCs). **b**, Three example RGCs from the purple cluster in **a**: Each sextet of PSTHs exhibits a delayed response to positive flows compared to negative flows and gratings (stimuli follow the same arrangement as in Fig. 1d). **c–e**, Example individual RGC responses to the six stimuli from other locations on the manifold indicated by lines; they exhibit distinct response profiles, e.g., no response to negative flows (**c**) or direction-tuned (**e**). **f–i**, The spatial receptive fields from each group of colored points form a mosaic-like arrangement that tiles space, confirming their identification as members of single classes of RGCs. *Left* panels show the 1-SD contour to a two-dimensional Gaussian fit to the spatial receptive fields estimated by computing the spike-triggered average to a checkerboard stimulus: colors correspond to the points on the manifold in **a**, hexagon shows the outline of the electrode array. *Middle* panels show the temporal receptive fields from the spike-triggered average from the same cells in the left panels. *Right* panels show an example sextet of PSTHs from a neuron in each mosaic to compare how the responses to the different stimuli vary across the different RGC types.

The observation of a largely discontinuous (or clustered) RGC encoding manifold is reminiscent of RGCs being organized into many distinct types [36, 81, 7, 41]. RGCs with nearly identical response properties are clustered on the manifold; those with distinct responses are segregated. Each of these RGC types exhibits highly stereotyped responses and feature selectivity while forming a mosaic of receptive fields that approximately tiles space [96, 30, 36]. To test whether clusters of cells on the encoding manifold corresponded to individual RGC types, we examined the spatial receptive field locations of RGCs within each cluster and from individual retinas by calculating the spike-triggered average to a checkerboard noise stimulus [19]; this revealed that RGCs in a given cluster, when sampled by the MEA at sufficient density, exhibited a mosaic-like organization (Figs. 3f–i and S4). This is a hallmark of individual cell types in the retina, indicating that the manifold embedding uncovered distinct RGC types. This observation serves further to validate, beyond the artificial ring model, the utility of the encoding manifold to reveal structure in a neural population.

We next applied the encoding manifold approach to MEA recordings from 640 mouse V1 neurons (Fig. 4 and S5d). The encoding manifold of V1 was quite different from that derived from the retina. Instead of clusters separated by abrupt changes, the V1 neurons were distributed relatively uniformly and continuously across the manifold (see Discussion). To emphasize this difference, for the V1 encoding manifold we speak of “neighborhoods” of cells, rather than distinct clusters, since preferred stimulus/response characteristics now vary smoothly from neighborhood to neighborhood.

**Fig. 4:**
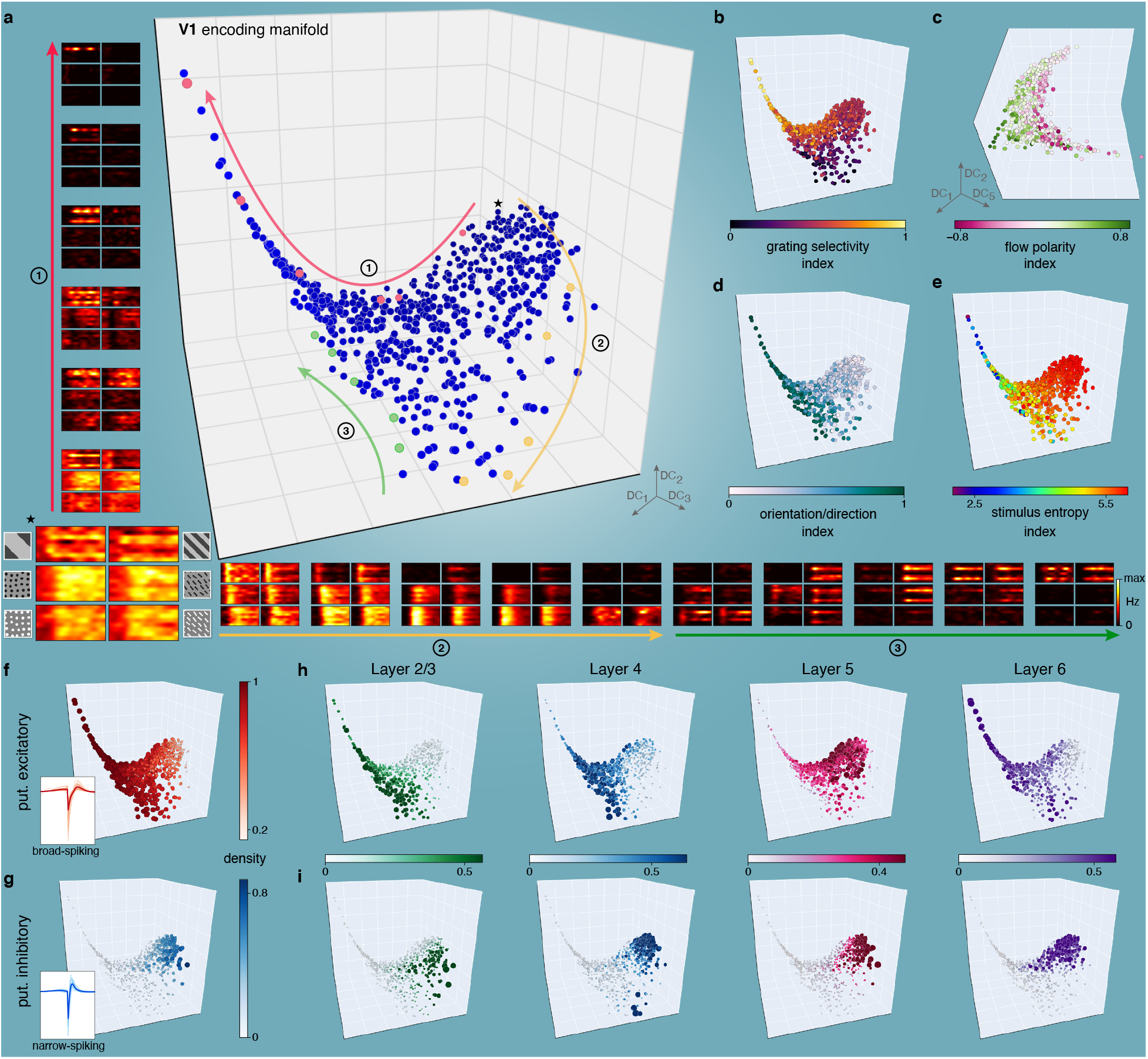
The encoding manifold formed by V1 neurons arranges neurons by their stimulus selectivity and response dynamics and is continuous rather than clustered. **a**, Projection of the encoding manifold formed by V1 neurons onto the first three diffusion coordinates (*n* = 640 neurons). The manifold consists of an extended arm formed by neurons tuned to low spatial frequencies connected to a higher-dimensional core of richly interconnected neurons. Instead of identifying clusters (as in Fig. 3), we follow paths over the manifold. Starting from the neuron indicated with *\**, along path 1 (red arrow) cells transition from responding to all stimuli, to being highly selective for gratings moving in a particular direction (response profiles shown correspond to neurons highlighted in red). Along path 2 (yellow arrow, yellow neurons), neurons transition from responding to all stimuli to preference for positive-contrast flows. Along path 3 (green arrow, green neurons), neurons that are highly orientation selective to flows transition to neurons that respond to gratings of both low and high spatial frequency. Throughout, the transition is gradual, indicating a continuous manifold. **b–e**, Visual feature selectivity (see Methods) is well organized over the manifold; different regions correlate with particular features, but there are no apparent gaps. **b**, Preference for gratings is highest along the arm. **c**, A higher diffusion coordinate reveals that contrast polarity preference varies smoothly. **d**, Orientation/direction selectivity declines toward the upper right. **e**, Distributed stimulus selectivity increases toward the upper right. **f–g**, Classification of neurons as putative excitatory or inhibitory (based on spike waveforms, see Methods) shows that physiological properties are also well-organized on the manifold. Different regions exhibit laminar specificity for both putative excitatory (**h**) and putative inhibitory (**i**) neurons, correlating with particular visual features from **b–e**. Neurons in **f**–**i** have color and size proportional to the local density of like-types in the data graph (see Methods).

While there are global coordinates (particular diffusion dimensions) that organize, e.g., contrast (Fig. 4c), a deeper picture emerges by following paths over the manifold. For example, Path 1 starts in a neighborhood of cells that respond to all stimuli before traversing an “arm” of cells selective only for low spatial frequency gratings (Fig. 4b). The origin of this path is a neighborhood populated primarily by fast-spiking putative inhibitory neurons (Fig. 4g), consistent with the observation that inhibitory neurons are broadly tuned [49]. Excitatory neurons in layers 2/3 versus layer 5 predominate in different regions on the manifold (Fig. 4h), consistent with the observations of mutual antagonism between these layers [73]. For other aspects of how the manifold organization relates to neuroanatomy and physiology, see Figs. S6 and S7.

The low spatial frequency arm is special; it consists mainly of excitatory neurons in layers 4 and 6, both of which, along with other layers, receive input from the lateral geniculate nucleus of the thalamus in the mouse [4]. Because these cells are known to be tuned to orientation or direction of motion, we can ignore this feature, making the manifold insensitive to their actual direction preferences. This approach allows us to focus on other, less explored, properties. For that reason, we factored out direction preference with a permuted tensor factorization (see Methods). This can readily be undone: when we isolated V1 neurons that responded primarily to low frequency gratings and that were orientation tuned, our embedding approach recovered a ring-like organization (Fig. S8), reminiscent of the ring model of orientation tuning (Fig. 2f). This illustrates that we are sampling a wide assortment of orientation-tuned neurons in V1, and that they organize with similar orientations supporting each other. However, we emphasize that these neurons accounted for only 35% of the population, and only 17% of the population had low frequency gratings as the sole stimulus eliciting a statistically significant response (see Methods).

The results above indicate that retinal output samples stimulus features in a relatively discrete manner, while V1 is substantially more continuous (Fig. S10b,c). This implies that cell diversity in the two circuits have conceptually different roles. In retina, each RGC type produces a distinct selectivity for a constellation of visual features. In cortex, cells appear to produce a continuum of stimulus selectivity that is relatively uniform across stimulus and response space, as one might expect either from a highly interconnected network or from a complete mixing of the parallel inputs from retina. We discuss implications of, and caveats to, this interpretation below.

## Encoding manifold for convolutional neural networks

Recently, convolutional neural networks (CNNs) have been widely used to model visual processing in retina and cortex. While they recapitulate at least some features of visual processing, such as a hierarchical architecture that explicitly represents progressively more complex features in visual scenes, they also lack the dense recurrent connectivity present in most neural circuits [99, 98, 58, 76], [11, 57, 10]. If CNNs were to accurately represent the way that populations of neurons encode visual features [82], the structure of their encoding manifolds should be similar to those found in biology. To test this, we constructed encoding manifolds from units in two popular deep CNNs: ResNet50 [45] (Figs. 5 and S9) and VGG16 [88] (Fig. S10d). Since these networks were trained on ImageNet [29], we first established that our stimuli were classifiable, and that they exhibited comparable activation levels across layers (Fig. 5b). Unlike V1, the encoding manifolds produced from CNNs were remarkably discrete (clustered) in their organization, even more so than the retina. For CNNs, most clusters corresponded to a feature map (set of units spanning position and sharing identical weights), reminiscent of RGC mosaics (Fig. 5e), revealing that activity patterns across feature maps were largely uncorrelated. These results indicate that CNNs do not encode visual feature space as neurons in V1 (and presumably other cortical areas) do, suggesting a crucial limitation to the use of CNNs for understanding cortical function (see Discussion).

**Fig. 5:**
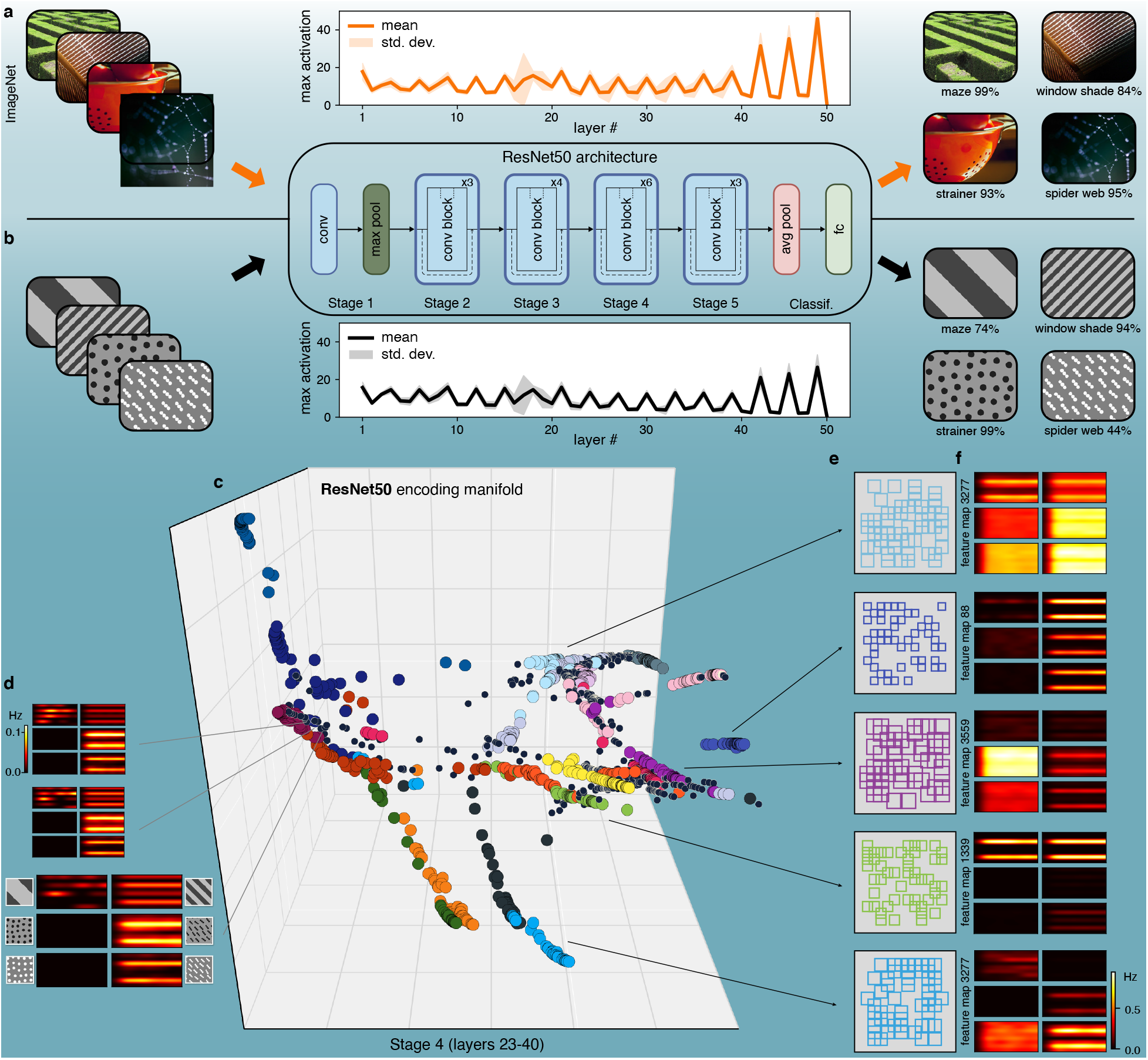
Encoding manifold of deep convolutional networks. **a**, ResNet50 is a popular CNN pre-trained on ImageNet for image classification. **b**, When applied to ResNet50, our stimuli (Fig. 1) generate similar levels of activation as those from natural scenes, and are also classified with confidence. **c**, Encoding manifold computed from neuronal units from Stage 4, sampled at random in proportion to their activation (*n* = 2000, see Methods; other stages are shown in Fig. S9). Color labels identify feature maps, showing that most neurons that share weights are grouped into well-defined clusters. **d**, Although most individual neurons exhibit response profiles similar to those from V1, the topology of the manifold obtained is strikingly different (compare with Fig. 4). **e**, Five examples of the spatial receptive fields (small squares) and response profile centroids of groups of neurons belonging to the same feature map. These form clusters that tile visual space (cf. RGC types, Fig. 3) and have very specific feature selectivity, as expressed by their distinct response profiles.

## Discussion

We began with a seemingly simple question: whether populations of RGCs and cortical neurons sample visual space similarly. We developed a novel, data-driven approach for topologically organizing how neurons in a circuit represent or encode stimuli by producing an “encoding manifold.” In this manifold, each point is a neuron and those with similar selectivity and response dynamics are nearby on it (Fig. 2). Remarkable topological differences emerged between retina and V1: The retinal manifold was clustered (Fig. 3) while the cortical manifold was much more continuous (Fig. 4). When comparing these manifolds to that of CNNs, popular models of cortical visual processing, we found that CNNs exhibited a topology quite distinct from that of V1 and more clustered than that of the retina (Fig. 5). A density-based hierarchical clustering algorithm (see Methods) confirms this qualitative assessment (Fig. S10a–c).

### Differing functional consequences of cell type diversity in retina and V1

Experiments beginning with Kuffler, Hubel, and Wiesel have demonstrated important differences in the responses of retina and V1 [53, 47]. However, those investigations have largely focused on differences in the optimal features that drive the cells (e.g., orientation tuning). Our purpose was distinct: to identify how stimulus space is sampled across each neural population, and to develop a method to visualize and quantify these differences. As such, we have not focused on the stimuli that generate peak responses, but instead we have used the response dynamics of each neuron to a battery of stimuli to learn how the population collectively samples stimulus space. For example, it is possible that retina and V1, while being sensitive to different visual features, could organize the encoding of these features similarly and thereby exhibit similarly clustered (or continuous) encoding manifolds. Furthermore, in retina and V1, a revolution has recently occurred in our understanding of cell type diversity, driven by connectomics and transcriptomics [86, 41, 9, 85, 101, 94]. In both retina and V1 there appear to be a large number of genuinely distinct cell types, rather than a continuum of morphologic and/or transcriptomic profiles. In retina, it is clear these transcriptomic and morphologic cell types correspond (at least) nearly one-to-one with functionally distinct types [41]. Our analysis recapitulates a “clustered” view in the retina, but produces a very different view of V1. Despite cell type diversity in both structures, the encoding manifold reveals a nearly continuous sampling of visual features in V1, while it produces a more clustered or discrete sampling in retina. In the retina, the elongated clusters reveal only modest quantitative changes in the relative magnitudes of responses to the different stimuli, without changing the selectivity (Fig. S4). In V1, on the other hand, there are several dimensions through which stimulus selectivity and response dynamics vary qualitatively (Fig. S6 and Methods, Sec. 1.8).

A potential caveat to the above conclusions is the choice of visual stimuli. We used stimuli spanning spatial frequency, contrast polarity, direction of motion, and orientation. Except for temporal frequency, this accounts for possibly the most commonly explored features in the vision literature. There are potentially a huge range of visual stimuli (*N_s_*) to which one might measure responses, and of course, there are a large number of neurons that might respond variously to the different stimuli (*N_n_*). In principle, there might be *N_s_ N_n_* distinct response types, and because responses are extended in time this number is much larger. The encoding manifold is a simplified representation of this huge range of potential variation. To the extent that trajectories across the manifold correspond to distinct properties of visual stimulation, properties that we can make sense of, the manifold embodies an understanding of the real range of variation encoded in the neural circuit. This, combined with the fact that the stimuli we used also produced strong responses from artificial networks trained on natural scenes (Fig. 5b), leads us to believe that the topological differences we found should remain, even with richer stimulus ensembles. Nevertheless, expanding the stimulus set is an important direction for future research. One challenge is to produce stimuli that are at least approximately isotropic in space like flow stimuli (to mitigate the impact of cells having receptive fields at different retinotopic locations), yet retain naturalist structure [35]. Features that could be added to our stimulus ensemble in a relatively straightforward way are (1) flow stimuli that temporally vary in their contrast or mean luminance, (2) superimposing flow stimuli moving at different speeds to mimic depth and parallax when animals move through natural scenes, and (3) including chromatic content. These stimulus variations would allow exploring how populations of neurons are organized with respect to contrast and luminance adaptation, depth, and chromatic tuning, all of which are exciting avenues for future work.

### Relationship between manifolds and neural circuits

We have established the relationship between encoding manifolds and the response properties of neurons to multiple stimuli via the construction of the data graph from our similarity kernel (Fig. 2c). While this can be viewed as a kind of abstract circuit, we emphasize it is not expected be one-to-one with the anatomical circuit. The manifold represents neural activity, i.e. the responses to the stimulus ensemble; the relationship between such responses and anatomical connectivity remains complex. For example, when neurons are nearby on the encoding manifold, it may be because those neurons receive common input (i.e., RGCs of the same type) or because they are synaptically connected to neurons with similar tuning and dynamics (i.e., the ring model), or both. Nevertheless, some circuit-level inferences do arise from the encoding manifolds. For example, spectral graph theory [90] suggests that the topological differences between retina and V1 are likely produced by differences in their global organization: the retina has relatively fewer connections (RGCs receive common input but are not generally synaptically coupled except by relatively weak gap junctions), while V1 is much more densely interconnected with recurrence across many scales [61], particularly among excitatory neurons with similar tuning [50]. To understand this, consider the stochastic block model [46, 1] from social networks, a random model in which edges are more likely to be drawn between nodes belonging to the same “community”—or circuit (Fig. 6). If circuits are completely disjoint, the *adjacency matrix* (dimensions of which are neurons neurons) has a block-diagonal structure: units within a block are connected; those in different blocks are not (see Methods). The resulting manifold embedding then consists of completely disconnected components, or clusters, as in most of our CNN examples (Figs. 5, S9, S10d). As connections begin to couple these circuits, the manifold embedding begins to join components, until the connections become sufficiently numerous to mimic the more continuous V1 manifold. We should note that the stochastic block model perspective is only a start for guiding intuition; it is limited because it explicitly assumes the presence of blocks with uniform connection probability [79].

**Fig. 6:**
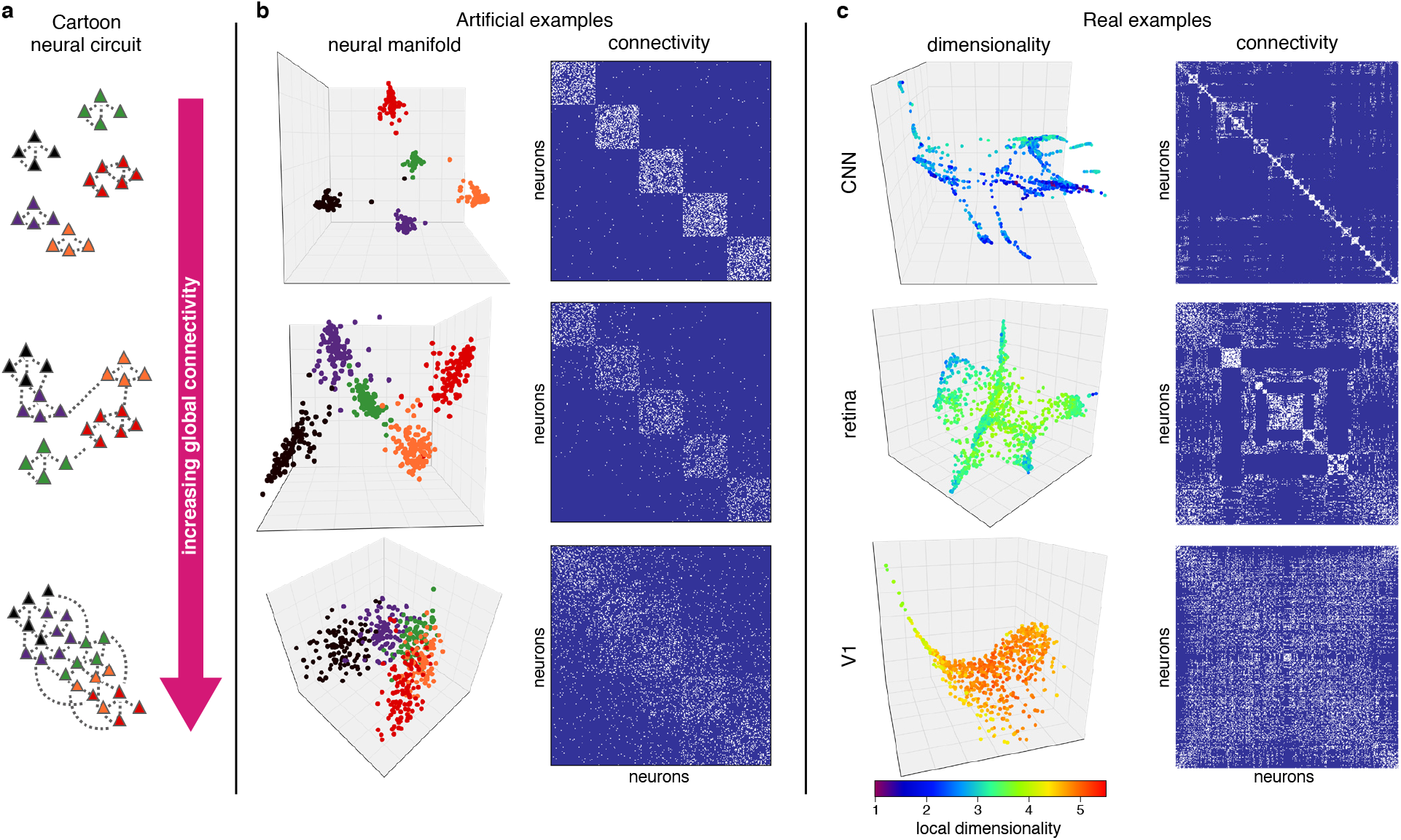
Circuits, connectivity, and dimensionality. **a**, Three examples of cartoon neural circuits with increasing connectivity. **b**, Extending the cartoon examples to artificial (stochastic block) models illustrates the relationship between the connectivity of the data graph and the resulting manifold. Neuron colors identify different blocks (left); connectivity is represented by adjacency matrices with connections in white (right). When dominated by intrablock connections, the manifold is disconnected (top). As connections between blocks are added, the manifold becomes more continuous (bottom). **c**, Adjacency matrices for the data graphs producing the manifolds from Figs. 3–5 range from diagonally-dominant (top) to a more widespread connectivity (bottom). V1 is continuous; retina has clusters with some overlap; and deep convolutional networks are even more clustered. The manifolds are now colored by their local intrinsic dimensionality, which also increases from top to bottom, underlining how network complexity is higher in cortex compared to the retina and CNNs (see Methods, section 1.8).

### Comparison with deep networks

Recently there has been great interest in assessing how artificial deep networks compare to biological networks [24]. This interest is motivated by the ability of CNNs to mimic many features of, for example, hierarchical visual processing found in macaques [99, 98, 76, 11, 58, 10, 57, 89]. However, the process of building an encoding manifold revealed a very important difference between CNNs and V1. While individual units in a CNN may exhibit tuning like typical cortical cells, their encoding topology is completely distinct from a population perspective: instead of being nearly continuous like V1, they are highly clustered, where each cluster typically identifies with a feature map (akin to RGC mosaics). Surprisingly, even feature maps across successive layers are largely uncorrelated in their activity patterns, which explains the lack of continuity in the CNN encoding manifolds (Figs. S9 and S10d). The encoding manifold can be used in future studies for directly testing how modifications to artificial networks, such as recurrence, affect their encoding topology and similarity to V1 and/or higher visual areas.

## Conclusion

In 1962, Hubel and Wiesel [47] noted that two novel response properties, not present in its inputs from the retina and dLGN, emerged in the visual cortex of the cat: orientation selectivity and binocularity. These properties were thought to be the product of the cortical circuit in combining the inputs it received. The absolute novelty of visual cortical responses is less clear in the mouse, with some degree of orientation selectivity and possibly binocularity in its inputs [93]. However, the encoding manifold of mouse V1 reveals that, rather than preserving the qualitative distinctions evident in RGC responses, the cortex combines its parallel inputs to create a novel mode of organization: carpeting stimulus space, rather than discrete sampling.

The best way to compare visual areas within species, across species, or with artificial networks has been an important question. Our encoding manifold introduces a new way to do this by providing a global (as opposed to pairwise [6]) view of how populations of neurons are organized to encode diverse stimuli. As we showed, it is applicable across areas and species. Furthermore, labeling the neurons on the manifold with known anatomical properties permits inferences about how physiology correlates with visual feature selectivity. Crucially, it facilitates the leap from abstract manifolds into actual biological circuits; see Fig. S7. Next questions implied by our results concern the dLGN and whether it is topologically more like the retina or the cortex, as well as extrastriate areas, for which there may be important differences in dimensionality and geometric properties. We believe this approach will also be useful for analyzing gene networks and the structure of other high-dimensional data sets.

## 1 Methods

### 1.1 Retina experiments

#### Animal procedures

All retinal experiments were approved by the Duke University Animal Care and Use Committee. Adult mice (2-6 months, C57Bl/6J, Jackson Laboratories, 000664) of both sexes were used. Animals were kept on a 12-hour light/dark cycle with *ad lib* access to food and water. Prior to use, animals were dark-adapted overnight by placing the animal in a light-shielded box fitted with an air pump for circulation. All dissection procedures the day of the experiment were carried out in complete darkness using infrared converters and cameras. Mice were decapitated, eyes enucleated and placed into oxygenated room-temperature Ames solution (Sigma, A1420) during retinal dissection and vitrectomy as described previously [100].

#### Multi-electrode array recordings

A 1.5 1.5 mm piece from dorsal retina was placed RGC side down on a MEA of 519 electrodes with 30 *µ*m spacing [38, 37, 74].

Oxygenated Ames solution perfused the retina throughout the experiment at a rate of 6–8 mL/min, heated to 32*^◦^*C. Hexagonal retinal recording array size in Figs. 3 and S4 is 0.49 mm across and subtends about 15 degrees of visual angle.

#### Spike sorting

Raw voltage traces from the MEA were spike sorted using YASS followed by manual curation [56, 55]. Briefly, spikes were identified by events that crossed a threshold set to 4 standard deviations from the mean voltage. The electrical event 0.5 ms preceding and 1.5 ms following this threshold was extracted from the recording. These events were accumulated on each electrode. Projection pursuit was used to to reduce the dimensionality of these signals and identify clear clusters of spikes. Putative cells with a spike rate *>*0.1 Hz and with *<*10% contamination estimated from refractory period violations were retained for further analysis.

#### Visual stimuli and receptive field measurements

The image from a gamma-calibrated OLED display (Emagin, SVGA + XL Rev3) was focused onto photoreceptors using an inverted microscope (Nikon, Ti-E) and 4x objective (Nikon, CFI Super Fluor x4). Checkerboard stimuli were created and presented using custom Matlab code and presented for 30 min to estimate spatial and temporal receptive fields. The checkerboard stimuli were presented at a photopic light level (about 10,000 Rh*/rod/s), and a new checkerboard pattern was presented every 33 ms. Each square in the checkerboard stimulus was 75×75 *µ*m. Custom software was used to present drifting gratings and flow stimuli at 60 Hz refresh rate, described in section 1.3 below. RGC responses to checkerboard noise were used to estimate the spatial and temporal components of the spike-triggered average (STA) [19]. The STA estimates the spatial and temporal integration of visual stimuli by the receptive field. Spatial receptive fields were fit with a two-dimensional Gaussian function to estimate the spatial extent of the receptive field center. A 1-standard deviation contour of this fit was used to summarize the size and location of receptive fields of each RGC. The mean intensity of the grating and flow stimuli was also about 10,000 Rh*/rod/s.

#### RGC classification

The time course of the temporal receptive fields, the autocorrelation function of the spiking dynamics and spatial receptive field size information were used to classify RGCs into different types, as described previously [74]. Distinct functional types were confirmed by the formation of receptive field “mosaics” [31, 36]: receptive fields that uniformly tile space and overlap at their 1-standard deviation contour.

### 1.2 Cortex experiments

#### Animal procedures

Experiments were performed on adult C57BL/6J mice (age 2–6 months) of either sex. All protocols and procedures are approved by the University of California–San Francisco Institutional Animal Care and Use Committee. Animals were maintained on a 12 h light/12 h dark cycle. Recordings were performed during the dark, more active phase of the cycle.

#### Preparation of mice for extracellular recording on the spherical treadmill

Recordings were done on alert mice free to run on a polystyrene ball (200 mm diameter, Graham Sweet Studios) floating on an air stream from a single inlet at the bottom of a hemispherical bowl of slightly greater inside diameter. During recordings, the animal’s head is fixed to a rigid crossbar above the floating ball by screwing a titanium or stainless steel headplate cemented to animal’s skull before recording using surgical procedures as described by [69]. Following recovery from surgery for headplate attachment, the animal is allowed to habituate to the recording setup and learn to control the ball.

#### Visual stimuli

Visual stimuli were presented with gamma-corrected video display (Nanao Flexscan, 30 × 40 cm, 60 Hz refresh rate, 32 cd/m^2^ mean luminance, or Dell Ultrasharp 38 cd/m^2^ mean luminance) placed 25 cm from the mouse, subtending 60*^◦^*–75*^◦^* of visual space.

For current source density (CSD) analysis, we present a contrast-reversing square checkerboard (0.04 cpd, squarewave reversing at 1 Hz). Other visual stimuli were the same as those described for the retina experiments, and are described in section 1.3 below. All stimuli variations were repeated 20–25 times according to a randomized sequence.

#### Extracellular recording in awake mice

To carry out microelectrode recordings, a craniotomy was performed under brief isoflurane anesthesia, and the skull was thinned over a 1–2 mm diameter centered above the monocular zone of V1 (2.5–3 mm lateral to midline, 1–2 mm anterior to lambda). At least 1 h after full recovery from anesthesia, this small opening allowed insertion of a 1.1-mm-long double-shank 128-channel probe [33], fabricated by the Masmanidis laboratory through the NSF NeuroNEX program (University of California–Los Angeles). The electrode was placed at an angle of 30*^◦^*–45*^◦^* to the cortical surface and inserted to a depth of 500–1000 *µ*m below the cortical surface. An additional period of 30 min to 1 h was allowed to pass before recording began. For each animal, the electrode was inserted no more than twice. Microelectrode and stimulus synchronization data were acquired using an Intan Technologies RHD2000Series.

#### Single-neuron analysis

Single units in earlier experiments were identified using MountainSort [21], or in later experiments Kilosort 3 [71], in both cases followed by manual curation. For a few experiments in which the raw data that had been sorted with Mountainsort were later sorted with Kilosort, Kilosort found more than 90% of the same units plus 10–60% of additional well-isolated units. Data from 323 units from 5 experiments in 3 mice of the [35] report were combined with new data from 317 units from 12 experiments in 12 mice to create the dataset used in the present report. Individual neurons were classified into broad spiking (putative excitatory) or narrow spiking (putative inhibitory) based on their extracellular spike waveform (see [68]).

#### Cortical layer

The cortical layer containing each isolated unit was determined using current source density (CSD) analysis on data collected during presentations of contrast-reversing square checkerboard. Briefly, extracellular voltages sampled at 20 kHz are bandpass filtered between 1 and 300 Hz to obtain local field potentials (LFPs) and then averaged across all 1 s positive-phase presentations of the checkerboard. Second spatial derivative of the average LFP traces along the length of the silicon probe provides us with the profile of CSD. The borders between layers 2/3–4, 4–5, and 5–6 are identified by spatiotemporal patterns of sinks and sources in the CSD plot (for example see Fig. 1c of [28]).

### 1.3 Visual stimuli

To characterize neural responses with single-unit recordings, we presented interleaved drifting square-wave grating stimuli and flow stimuli moving in 8 directions at a temporal frequency of 4 cycle/s and 50% contrast, with a trial duration of 1.25 s. Spatial frequencies used for gratings included 0.04, 0.15, 0.24, and 0.5 cycle/deg. As in [35], we used flow stimuli with two different geometries. The first were non-oriented single-dot flows, and the other were oriented flow elements with 3 collinear dots. Both oriented and non-oriented variations had one version with positive contrast (white dots against a gray background), and another with negative contrast (black dots against a gray background). Dominant spatial frequency contents of 0.15, 0.24, and 0.5 cycle/deg were used, corresponding to the following dot diameters, in degrees of visual angle (respectively, approximate dot spacings, in multiples of diameter): 2.1 (2), 2 (1), 1 (1) for single dots; for 3-dot flows, diameter was divided by 3 to preserve the total area of each flow element.

Because the flow stimuli were stochastic (in position and velocity), we presented at least 3 different instances of each variation, which were repeated to account for the desired number of total trials (at least 10 for retina, and 20 for V1).

### 1.4 Embeddings of trials for decoding manifolds

To produce the decoding embeddings (Fig. 1f), we ran principal component analysis (PCA) using the implementation of the `scikit-learn` Python package [72] on a matrix whose dimensions were neurons by total number of trials. Each entry consisted on the average firing rate for that trial.

### 1.5 Ring model

The ring model of [13] is a model of an orientation hypercolumn, inspired by the primary visual cortex of cats and monkeys [48]. It is used here as an example to illustrate the development of our approach (see Fig. 2). Neurons are parameterized by their preferred orientation (PO), *θ*, an angle between *π/*2 and *π/*2. Both excitatory and inhibitory neurons (*n* = 250) form recurrent connections to all others, with the strength of interaction depending on the difference between their POs.

The net interaction between neurons *θ* and *θ^′^* is given by *J*(*θ θ^′^*) = *J*_0_ + *J*_2_cos(2(*θ − θ^′^*)), where *J*_0_ = 80 is the amount of uniform inhibition, and *J*_2_ = 112 the orientation-specific part of the interaction (see connection weight matrix in Fig. S1). The activity profile of the network at time *t* is given by

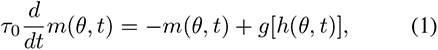

where *τ*_0_ = 5 ms is a time constant, and *g* a semilinear gain function with threshold *T* = 1 and gain factor *β* = 0.1 (refer to [13] for details). All neurons receive external input expressed as *h*^ext^(*θ*) = *c*[1 − *ɛ* + *ɛ*cos(2*θ*)], where *ϵ* = 0.5 regulates its angular anisotropy and the coefficient *c* = 1.9 sets the stimulus contrast. The total synaptic input at time *t* of a neuron *θ* is

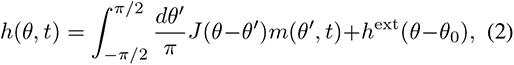

where *θ*_0_ is the orientation of the external stimulus.

After receiving an oriented input (the external stimulus), each artificial neuron responds proportionally to the similarity between the orientation of the stimulus and its own preference. Its ultimate change in output (treated as spikes/s) is obtained by integrating the inputs from all its neighbors plus the external input. In our simulation, 8 different stimuli were used with orientations evenly distributed between *π/*2 and *π/*2, and a trial length of 1.25 s. Each stimulus was presented 10 times, with a Gaussian noise stimulus (std. dev. 3) used during the ISI to make the response in each trial essentially unique, given that the model is deterministic.

### 1.6 Data analysis: tensor decomposition

This section is organized as follows. We begin with several essential points about tensor factorization (Fig. 2b), then specify our tensor explicitly. For general background, see [97]. Next, we show how the neural factor matrix can be used to define the neural encoding space (Fig. 2c), and address the question of how many tensor factors to use. These are essential steps that will allow us to define similarities between neurons and infer the neural encoding manifold (Fig. 2).

#### Tensor decomposition

In a tensor CP decomposition [18, 44], an *n*-way tensor ***T*** R*^I_1_×I_2_×…×I_n_^* is approximated by a sum of rank-1 tensors. Let *R* be the number of components chosen; then, following the notation of [51], we may express this for the specific case of a 3-way tensor as:

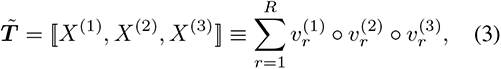

where *◦* stands for the vector outer product and *X*^(^*^k^*^)^ is called a *factor matrix* containing the factors *v_r_*^(^*^k^*^)^ as its columns. Each rank-1 tensor is formed by the outer product between each factor in the same component (following the convention that a *component* refers to each set of associated factors, one from each tensor mode).

It is often convenient to normalize factors to unit length and collect their original magnitudes into a single scalar for each component, 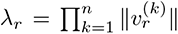, where *r* indexes the component and *k* the factor mode, for an *n*- way tensor. Combining all scalar into a vector *λ* ∈ R*^R^*, we may rewrite eq. 3 as

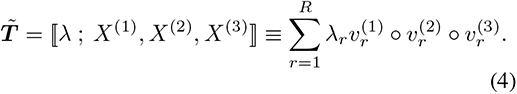

Most algorithms for CP decomposition use the squared reconstruction error as objective function [23]:

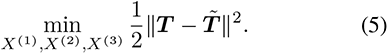

where is the norm of the vectorized tensor (analogous to the Frobenius norm for matrices). Since the data consists of neuronal firing rates, we compute a non-negative tensor decomposition, also known as non-negative tensor factorization (NTF) [15], formulated as in eq. 5 with the additional non-negativity constraint *X*^(^*^k^*^)^ 0*, k*. In contrast to the negative version, it achieves a parts-based representation with more easily-interpretable components [54, 23]; moreover, it can produce more stable factors, is less likely to overfit, and has comparable parameter efficiency [97]. To compute the decomposition, we use the gradient-based direct optimization approach (OPT) [2] via the Tensor Toolbox implementation [8], adapted to allow permutation of the response factors (see below).

Items to note: the factor matrices are not orthogonal and may in fact have linearly dependent columns. Importantly, the best rank-*r* factorization may not be part of the best rank-(*r* − 1) factorization [52]. Because tensor factorization requires an initial guess for the factors, each run may yield different results, even when the same number of components is chosen. Reconstruction error is commonly normalized by the Frobenius norm of the original tensor (e.g., [97]), and the values shown in Figs. S2d, S3c, S5c, S8e follow this convention.

#### Tensor of neuronal responses (Fig. 2b)

We utilize a 3-way tensor having neurons, stimuli, and responses as modes. We choose not to use individual trials since there is no expectation of significant change in the responses over different repetitions of the stimulus (in contrast with a learning task). Moreover, by using stimuli as a mode, we obtain more interpretable factors that directly relate to parts of the stimulus ensemble. For each neuron recorded, and stimulus used, we have a response, which could be either a 1-D response curve—as was the case for the ring model (Fig. S1) and the embedding of V1 using low frequency gratings only (Fig. S8)—or a 2-D response map (stimulus direction × time; see Fig. 1). The latter was used in vectorized form (concatenation of the responses to the 8 directions of motion), thus forming a 3-way tensor.

The responses of a neuron to each stimulus were kernel-smoothed using the Improved Sheather-Jones algorithm for automatic bandwidth selection [14], which does not assume normally distributed data; we used the `KDEpy` Python implementation[70]. For the bandwidth fitting, we combined all trials in which the neuron was responsive (at least 5 for retina, and 15 for V1). Kernel bandwidths were allowed to be automatically selected within the range 10–50 ms, using the stimulus direction with the highest number of spikes.

#### Neural encoding space (Fig. 2c)

Next we define a matrix formed by the neural factors obtained from NTF, and define the space associated with it, from the perspective of linear algebra. We refer to this as the neural encoding space (see Fig. 2c). From eq. 4, we can express the reconstruction of an individual data point, *x_i_*, as:

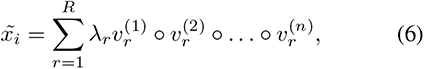

and for the entire tensor as:

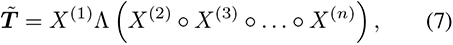

where Λ is the diagonal matrix with entries Λ*_rr_* = *λ_r_*.

Since the neural factors are represented by the first mode of the tensor, we henceforth denote *X*^(1)^ by . Now note that the outer product between the vectors of the remaining modes of each component is a (*n* − 1)-way tensor, which, if vectorized, becomes a unit-norm vector, and can be considered a basis vector. Technically, this collection of vectors (one for each component *r*) represents a *frame* [20], since in general it will be redundant (containing more vectors than the true dimensionality of the space that they span). We refer to it as the *stimulus-response frame*.

We can then define a matricized version of ***T*** with respect to the first mode, denoted by *X*_(*N*)_ *∈* R^(*I*_2_*×…×I_n_*)*×I*_1_^, as:

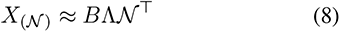

with each column *b*:*_,r_* of *B* given by

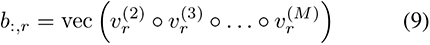

for *r* = 1*, …, R*. These *R* columns can therefore be regarded as unit-norm basis vectors. So each data point ***x****_i_* (column of *X*_(_*_N_* _)_) is approximated as a non-negative linear combination of the vectors in the stimulus-response frame.

In this case, it is convenient to multiply each column *v_r_*^(1)^ of *N* by its corresponding *λ_r_* and work with a scaled neural matrix, *N_λ_*:

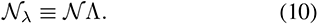

The matrix *N_λ_* thus contains all information needed to reconstruct each neuron *x_i_* in terms of the stimulus-response frame: its linear combination coefficients are given by the *i*^th^ row of the neural matrix *N_λ_*. The dimensionality of the data points is thus reduced to *R*. We now discuss the procedure for finding *R*.

#### Choosing the number of tensor components

The neural matrix depends on the number of components, *F*, a choice that is challenging and computationally costly [103]. The standard procedure would be to minimize reconstruction error, where one would use the number of components sufficient to meet a prespecified tolerance. Our goal is different, however: We seek to find those features that maximally organize the data. That is, we are concerned with the span of the latent space and the stability of neural matrix. We now develop this idea.

When *F* is too small, there are not sufficient factors to capture all the relevant structure present in the data. When it is too large, degeneracy of the resulting components may arise, a phenomenon known as overfactoring [2]: some factors could mostly capture noise, and otherwise meaningful factors could be split. Because in NTF the factors are in general not orthogonal, some degree of redundancy typically exists. Applying PCA to the neural matrix reveals its redundancy by re-expressing it in terms of an orthonormal basis formed by the eigenvectors of the sample covariance matrix. Its eigenvalues express the variance of the data when projected onto each eigenvector. Looking at this spectrum allows one to estimate the rank of, or the intrinsic dimensionality of the latent neural space. Intuitively, while more factors should explain more variance, at some point the gain can become insignificant or a product of overfactoring. By analogy with overfitting, we argue that the best *F* is the smallest one that maximizes the variance explained by the neurons in this low-dimensional subspace. As we now show, this choice is highly tractable.

An example factorization is that of the ring model from Fig. 2: each component is responsible for a single stimulus, and neurons overlap slightly as one moves from one component to the next (Fig. S1). This is predictable from the simulation setup: each neuron’s response function decreases slowly as a function of the difference between their preferred orientation and the orientation of the input stimulus. Now since all neurons respond with the same temporal dynamics (minus noise), only a single response pattern is required to reconstruct the responses to any of the 8 stimuli in the original tensor. Therefore *F* =8 is necessary and sufficient, regardless of the number of neurons used.

Intuitively, one might want *F* to maximize the span of the neural factors. However, since the factors are in general not orthogonal, this number might greatly exceed the intrinsic dimensionality of the data. When each of the neural factors has unit norm, the spectrum of its covariance matrix reflects how much redundancy exists in the frame, so instead of using the span directly, we use the nuclear norm (sum of eigenvalues, or variances) as a measure of the (putative) rank of . For a fair comparison across different choices of *F*, then, we normalize the variance spectrum by dividing by its maximum eigenvalue (since uniformly scaling the data points will not change the embedding). This approach is illustrated in Fig. S2 for the ring model. In particular, notice how the nuclear norm tends to increase rapidly for small *F*, then reach a point where it plateaus (Fig. S2d). For real data, we observe similar behavior (Figs. S3b, S5b, S7d, S8d, and S9b). We denote this point as *F* = *R*, representing a kind of structural rank for the neural matrix. As *F* increases further, the total variance tends to keep increasing albeit at a lower rate, since the resulting components either overfit or break down pre-existing factors [2].

One detail remains with regard to NTF, since a random initialization is used in the algorithm. We use 50 different initializations for each value of *F*, and choose the best-fit model (minimum reconstruction error) within *F* = *R*. An alternative is to use the average weighted graph from all different initializations; it produced similar results.

#### Permuted tensor decomposition

As indicated in the main text, normalizing the neurons’ preferred direction of motion allows the manifold inference to focus on other, less explored, properties. For example, two neurons narrowly-tuned to orientation can be considered to be similar regardless of their particular preferred orientation (in Fig. S8, we show how orientation or direction preferences can be considered explicitly). Thus for the retina, V1, and deep networks datasets we develop an algorithm in which the decomposition can automatically search through alternative versions of the data tensor to minimize reconstruction error. This allows one to solve for factors that are agnostic to the actual orientation/direction preference of each neuron. The direct optimization (OPT) method from [2] is well-suited for this because it solves the CP decomposition using gradient descent, solving for all factor matrices simultaneously (as opposed to traditional methods using alternating least squares [18, 44]). As shown below, this makes it feasible to incorporate permutations of entries along one of the tensor modes directly into the optimization loop. In the above datasets, because the stimuli drifted in 8 different directions, such a strategy amounts to computing each step of the decomposition as the argmin of all possible circular-rotations of the rows in each of the 2-D response maps in ***T*** . As described above, since the response maps are vectorized to become tensor fibers, the circular-rotation effectively becomes a permutation of their entries.

Following [2], the objective function of eq. 5 can be expressed as three summands:

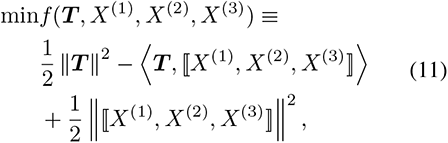

where the second summand is the inner product between ***T*** and its CP approximation.

At any given step *t* of the optimization, let its current factor matrices be expressed as [*X*^(1)^*, X*^(2)^*, X*^(3)^ ]^(^*^t^*^)^, and denote a permuted version of ***T*** as ***T*** *^′^*. To enforce the desired invariance, we shall replace ***T*** with the optimally permuted tensor, ***T*** *^*^*^(*t*)^, namely that obtained from the set of independent permutations of each individual response map in ***T*** resulting in the lowest value of *f* :

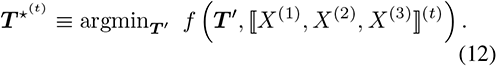

To find ***T****^*^*^(*t*)^, it suffices to compare the value obtained for the second summand in eq. 11 across all permutations (since the norm of ***T*** is unchanged by the permutations). Likewise, in computing *f* at step *t*, only that term is modified:

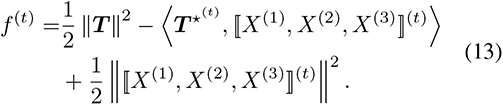

Finally, the corresponding gradient of *f* ^(^*^t^*^)^ can be computed by replacing ***T*** with ***T*** *^*^*^(*t*)^ in eq. 4.9 from [2]. The overhead of finding the optimal permutation is typically reasonable, because the tensor norm in the objective is computed elementwise. Thus the optimal permutation of each response map can be found independently. This results in an upper bound on scaling the computational cost of the objective by a constant factor, namely the number of permutations allowed (in the case of our stimuli, 8).

#### Fitting statistically non-significant responses

Responses to each stimulus were tested for significance by performing a two-tailed Mann-Whitney test comparing the mean activity during the second half of the ISI immediately preceding the stimulus versus mean activity over any interval of the same length within the stimulus presentation. This strategy allowed transient responses to be able to reach the same level of significance as sustained responses. Responses were deemed significant if they had *p <* 0.0005. When choosing between medium and high spatial frequency stimuli for each neuron, the optimal frequency was chosen as that which produced the largest number of significant stimuli among gratings, positive flows (1- and 3-dot), and negative flows (1- and 3- dot). Low spatial frequency gratings, because they had no flow counterpart at an equivalent frequency, were always included. See section 1.3 for more details.

Responses that were not statistically significant (see below) were removed (zeroed) from ***T*** before NTF to prevent introducing excessive noise. However, for the scheme to be able to reconstruct all data, after NTF we reintroduced these non-significant responses (see below). Responses that are truly non-significant are less reproducible and have very low magnitudes, so their neural coefficients will be close to zero. On the other hand, responses that are selective but too weak to pass the statistical test are likely to be well fitted regardless, since their response patterns will match those of other (significant) neurons. Moreover, using only highly significant responses in NTF ensured the factors obtained were interpretable and consistent.

#### Tensor normalization

We now discuss several details regarding the preprocessing of neural responses to construct the tensor. Since the objective of NTF is to minimize reconstruction error (eq. 5), response patterns with longer periods of activity (or with larger area over the response map) will have a higher reconstruction priority in terms of cost minimization compared to shorter responses. This creates a bias towards producing factors that reconstruct the sustained response patterns. We prevent this by pre-normalizing each stimulus response to have unit norm in ***T***.

Once the factorization is computed, the relative activity levels of a neuron to each stimulus can be reintroduced by rescaling the unit-norm responses by their original relative magnitudes. This is done by re-fitting the neural coefficients in *_λ_* to the rescaled responses using linear least squares. The rescaled tensor used in this step includes the non-significant responses (see above).

After this step, we reach a concrete interpretation for the neural coefficients: when a stimulus factor *s_i_* includes a single stimulus, its corresponding neural factor *n_i_* encodes the relative magnitude of the response pattern *p_i_* for each neuron in the population. Finally, for this to hold also in cases in which the same *s_i_* contains multiple stimuli, we make the following rebalancing between *n_i_* and *s_i_*. Let *s_i_^max^* by the maximum value in *s_i_*. Then, replace *n_i_* with *n_i_^′^* = *s_i_^max^n_i_* and *s_i_* with *s_i_^′^* = *s_i_*/*s_i_^max^*. Additionally, we take their square-roots so that the magnitude of the coefficients for each stimulus (when they are separable) approximates its relative response magnitude. This matters since the neuron organization should reflect not only the response patterns but also their stimulus preference in terms of relative firing rate.

### 1.7 Data analysis: neural encoding manifolds

**Low-dimensional embeddings (Fig. 2c, right)** Manifold embeddings were obtained by first computing the IAN similarity kernel [34] from the pairwise distance matrix obtained from the PCA projection of the neural matrix as input (see above). We used the number of principal components necessary to explain at least 80% of the variance. IAN uses a multiscale Gaussian kernel where the individual scales (or neighborhood sizes) are automatically inferred based on the local geometry of the data; its default parameters were used throughout [34]. The converged weighted graph (similarity matrix) from IAN is then used as input to the diffusion maps algorithm [26, 25], a nonlinear spectral embedding method based on the eigendecomposition of the normalized graph Laplacian. Diffusion maps were preferred over other popular methods (e.g., [95, 64]) because it is a deterministic algorithm and tends to better preserve the original topology of the data, cf. examples in [34]. The standard parameters *α* = 1 (Laplace-Beltrami approximation) and diffusion time *t* = 1 were used. It produces a set of diffusion coordinates which can be used to embed the neurons in stimulus-response coordinates (see above). When data graphs were disconnected, a high number of coordinates were required to embed all components together. To aid visualization, in these cases, we projected the diffusion distances between points using classical multidimensional scaling.

**Layer and type densities and encoding similarity.** Laminar and putative type densities for each cortical cell (Fig. 4) were computed as the fraction of adjacent nodes in the non-weighted IAN graph belonging to the same putative type and layer. The neuron densities were then arranged into a vector for each layer. Encoding similarities were computed as the non-negative cosine similarity between non-negative z-scores of the density vectors for different types and layers (Fig. S7).

### 1.8 Data analysis: intrinsic dimensionality

Intrinsic dimensionality plays an important role in manifold learning as a means to describe the degrees of freedom within the volume occupied by the data in ambient space, and it has been applied to the decoding approach [91] (Fig. 1f). In an encoding manifold, intrinsic dimensionality can be interpreted as the number of different directions of progressive change in stimulus-response space. For example, in the ring model the manifold is one dimensional, because it has a single latent variable, namely orientation, even though it is originally within a much higher-dimensional ambient space (the number of stimuli times the number of time points). Therefore, starting from a given neuron, orientation is the only variable that characterizes change when moving along the manifold since, for this artificial data, dynamics are constant. In CNNs, these latent directions correspond largely to variations in response magnitudes due to position over the feature map. In the retina, directions signify variation in stimulus preference, but these are highly constrained (Fig. S4). Across most of the V1 manifold the response profiles vary smoothly in several different ways, corresponding to different directions over the manifold (Fig. S6). The sole exception to this is the region where neurons respond only to low spatial frequency gratings, where the intrinsic dimensionality (ID) is mostly due to phase and tuning width differences, and direction vs. orientation tuning. Importantly, responses can vary only in some directions, and not others. One example is the absence of neurons with high orientation selectivity to 3-dot flows with a particular contrast polarity preference.

Importantly, even though the actual ID may vary with the variety of the stimulus set used [40], we believe that the ID of these three networks reflects intrinsic properties of their circuits, and should therefore be robust to variations in the stimulus set. To estimate ID of the manifolds, we used the neighborhoods of the unweighted IAN data graph as input to the Neighborhood Correlation Dimension algorithm [34], which can estimate the dimensionality around each data point (shown in Fig. 6).

The adjacency matrices from Fig. 6 had the neurons sorted according to the dendrograms produced by the density-based hierarchical clustering algorithm HDB- SCAN [17], shown in Fig. S10a–c and produced using the `hdbscan` Python library [63]. Its main parameter is the number of nearest neighbors, *k*; to avoid having to choose an arbitrary value for *k*, we instead use the neighborhoods inferred by the IAN kernel (section 1.7).

### 1.9 Data analysis: stimulus selectivity

Selectivity indices for various visual features (Fig. 4b–e) were computed as follows. Let the maximum response (instantaneous mean firing rate) to any of the grating stimuli (flow stimuli, respectively) be denoted as *G*_max_ (*F*_max_, resp.). A *grating selectivity index* for a given neuron was computed as the ratio *G*_max_*/*(*G*_max_ + *F*_max_).

Let the relative response magnitude of a neuron to a given stimulus be defined as the ratio between its maximum response to that stimulus and its maximum response to any stimulus (therefore yielding a number between 0 and 1). Defining *P*_rel_ (*N*_rel_, resp.) as the relative magnitude of a neuron’s response to any positive contrast flow (negative contrast flow, resp.), a *flow polarity index* was computed as the different *P*_rel_ *− N*_rel_.

An orientation selectivity index (OSI) was defined as: (*R*pref *R*ortho)*/*(*R*pref + *R*ortho), where *R*pref = (*R*_peak dir_ + *R*_peak dir+_*_π_*) is the response (average firing rate) for the preferred orientation and *R*_ortho_ = (*R*_ortho dir_ + *R*_ortho dir+_*_π_*) is the response for the orientation orthogonal to the preferred one. A direction selectivity index (DSI) was defined as: (*R*_pref_ *R*_null_)*/*(*R*_pref_ + *R*_null_), where *R*_pref_ is the response (average firing rate) for the preferred orientation and *R*_null_ is the response for the null orientation (*π* rad apart from the preferred one). An orientation/direction selectivity index (ODSI) was defined as the maximum value between a neuron’s OSI and its DSI.

A stimulus entropy index was defined as the 2*^H^*, where *H* is the base-2 entropy of the vector containing the relative response magnitudes of a neuron to the 6 stimulus classes used in the experiments (divided by their sum). It therefore ranges between 0 (case in which the neuron responds to a single stimulus) and 6 (when it responds with uniform magnitude to all stimuli).

Cortical cells were classified as well-tuned to low-frequency gratings by fitting their direction tuning curves to a double Gaussian, following the fitting method of [62] and selecting those cells with small fitting error, followed by manual curation based on their ODSI values. Their preferred stimulus direction used to label the embeddings in Fig. S8a,b was computed as the argmax of the fitted tuning curved.

### 1.10 Stochastic block model

For the toy circuit examples of Fig. 6b, we ran simulations using the traditional stochastic block model [46, 1], which models a network as a partition of nodes into blocks, or clusters. Edges are established between pairs of nodes according to prespecified intra- and inter-block probabilities. Each of the three networks used 5 blocks with *n* = 100 nodes per block. The top row network was modeled using a uniform intracluster edge probability *P*_intra_ = 0.1 and intercluster probability *P*_inter_ = 0.001. The middle row network used *P*_intra_ = 0.05 and *P*_inter_ = [0.005, 0.001, 0.001, 0.0001] as the intercluster probabilities to the other four clusters. The bottom row network used *P*_intra_ = 0.05 and *P*_inter_ = [0.025, 0.01, 0.005, 0.005]. Adjacency matrices in Fig. 6b,c were subsampled using a maxpool operation to have comparable dimensions and facilitate visualization.

### 1.11 Deep convolutional networks

We used publicly available models of the ResNet [45] and VGG [88] network architectures, pre-trained on ImageNet [29]. Both were state-of-the-art when released, excelling in tasks like classification and localization (see, e.g., [3] for a review). The former is still widely used; it introduced residual connections, a breakthrough in deep learning which has since been applied to most architectures, even to the most recent transformer networks. ResNet and VGG are particularly relevant to this study because both have been frequently used to assess how well they perform as models for the visual system in mouse [16, 87, 66] and primate [83, 104].

For their inputs (stimuli), we used cropped frames from our stimulus ensemble movie (conforming to the required size of these networks, namely 224 × 224 pixels) and fed them sequentially to the networks in order to obtain temporal activity from individual neuronal units (artificial neurons). Because of the necessary cropping, all stimulus frames were uniformly scaled to 50% of their original size in order to increase the number of visual features present in the same frame.

Activity levels in response to each frame were taken as the output of each unit’s non-linearity. For the classification layer of ResNet50, which uses a linear layer followed directly by a softmax operation, we half-rectified the outputs (equivalent to a ReLU). All other layers from ResNet50 (and all from VGG16) used a ReLU as nonlinearity. The numbering of layers from ResNet50 used in Figs. 5 and S9b–e follows the direct path in the network.

In analogy to what was done with the biological networks, we combined the responses to multiple trials into a single response map. Because there are no actual spikes, instead of applying the smoothing kernel (see above) we simply used a smoothing window of size 3 frames (equivalent to a 50 ms bandwidth, the maximum kernel width we used for real neurons). Receptive fields (RFs) were computed following the procedure from [5]. We display RFs for units from several feature maps in Fig. 5e, uniformly scaled in size for clarity.

#### Sampling procedure

Each manifold was built by sampling neurons from all layers in the same stage. We first chose 40 feature maps at random, in proportion to their maximum stimulus activity, to prevent the final manifold from including, by chance, only non-responsive feature maps. To eliminate border effects, sampling was concentrated on a central region of the feature maps. 50 neurons from each map were then sampled in proportion to their maximum activity, for a total of 2000 neurons (same order of magnitude as the retina, for comparable visualization). Examples of which neuronal units were sampled from the same feature map can be seen in Fig. 5e; each receptive field (RF, see above) corresponds to a sampled unit whose position on the response map is at the center of its RF. Within the fully-connected layers, where neurons do not share weights, there is no concept of a feature map, so we selected 2000 neurons at random, again in proportion to their maximum responses.

## Data availability

The datasets analysed during the current study are available from the authors upon reasonable request.

## Code availability

Code for methods described in this paper is available at https://github.com/dyballa/NeuralEncodingManifolds.

## Acknowledgements

Supported by NIH grant 1R01EY031059, NSF Grant 1822598, and the Swartz Foundation. We thank Mario Dipoppa, Lindsey Glickfeld, and Massimo Scanziani for a critical reading.

## Author contributions

SWZ, LD, GDF, and MPS wrote the manuscript. LD and SWZ performed the mathematical analysis. LD composed the figures. AMR, MT, and GDF made and analyzed the retinal ganglion cell recordings. MSH and MPS made and analyzed the cortical recordings.

---

**Fig. S1:**
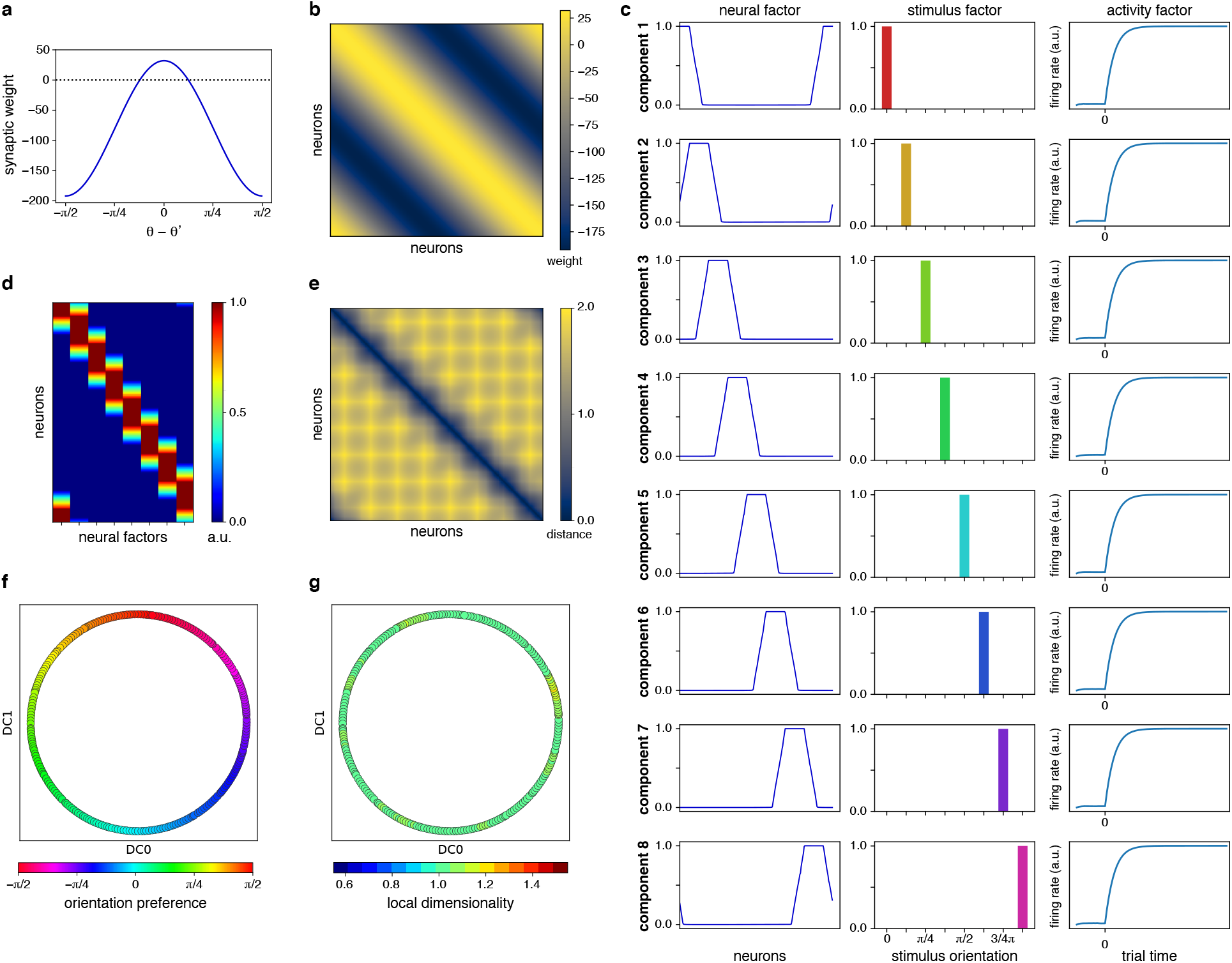
Encoding manifold algorithm applied to the ring model. **a**, Synaptic connection weights between pairs of neurons indexed by difference in preferred orientation. Only similar orientations support one another; otherwise are inhibitory. **b**, Adjacency matrix representation of synaptic interactions. Columns with similar orientation preference excite and those with different preference inhibit one another (see Methods, section 1.5). **c**, The 8 components obtained from non-negative tensor factorization (see Methods, section 1.6). Response dynamics were set to be nearly identical in this simulation and appear as such. Thus the components are dictated by the stimulus (orientation) preference. **d**, Neural matrix obtained by collecting the neural factors from **c** (see Methods, section 1.6). Each row represents the loadings of a neuron across neural factors. **e**, The pairwise distance matrix computed from **d** and used as input to the embedding algorithm. **f**, Diffusion map embedding using the adaptive neighborhood kernel (see Methods, section 1.7). **g**, The estimated local dimensionality (see Methods, section 1.8) is 1 throughout, compatible with the single latent variable in the encoding of the stimuli used, namely orientation.

**Fig. S2:**
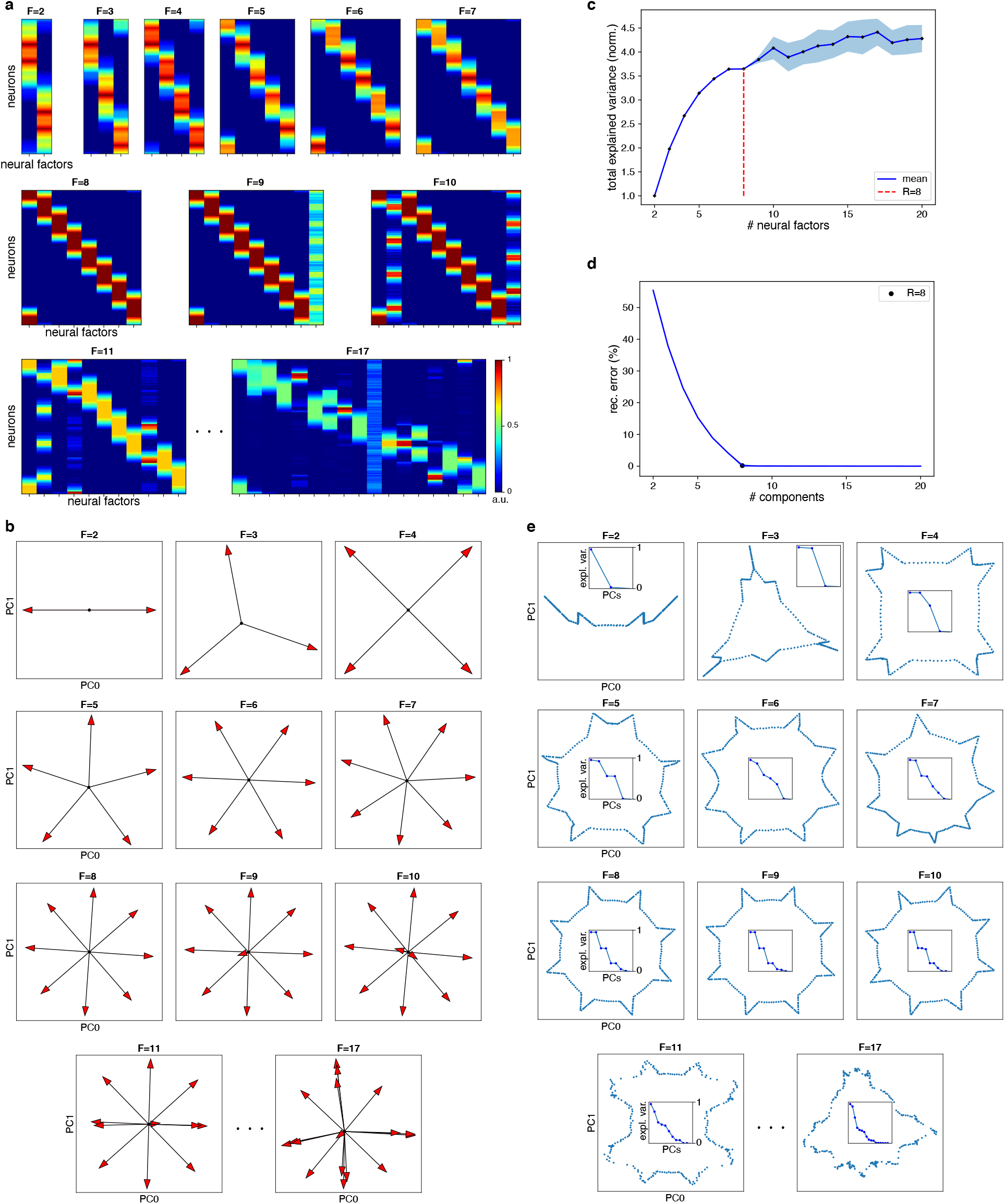
Choosing the number of tensor components, *F*, for the ring model. **a**, Neural matrices, denoted *N*, resulting from various values of *F* . The neural loadings are most uniformly distributed when *F* = 2, 4, or 8, by symmetry, but it is only *F* = 8 that maximizes the variance explained. When *F >* 8, observe the presence of factors that either reconstruct noise (therefore explaining very little additional variance), or split previous factors into two nearly identical vectors with loadings roughly halved. Both of these situations should be avoided (see Methods, section 1.6). **b**, An alternate representation in which the neural factors are plotted as vectors (arrows) projected onto the first two principal components of *N ^T^* . We now seek to maximize their span. Even though the ring can be embedded in a two-dimensional space, the volume defined by its neural factors finds a plateau only at *F* =8, since the factors are not orthogonal. This is precisely the quantity expressed by the (weighted) nuclear norm (see Methods, section 1.6). **c**, The total explained variance (nuclear norm) increases rapidly up to *F* =8, after which it flattens to a much lower rate. At this point, the variability across runs also increases markedly for larger *F* (cf. [97], their Fig. 2f). **d**, In this artificial example, reconstruction error alone (see Methods) is sufficient to decide the optimal choice of *F* = 8. **e**, Yet another representation: embeddings of neurons obtained by running PCA on the neural matrices from (**a**). The “ring” becomes maximally uniform at *F* = 8, and is stable until *F* = 10, above which the overfactoring makes it degenerate. Notice in particular the failure of a linear embedding algorithm to completely “unfold” the manifold (contrast these with Fig. S1f). Insets show the variance spectrum formed by the principal values.

**Fig. S3:**
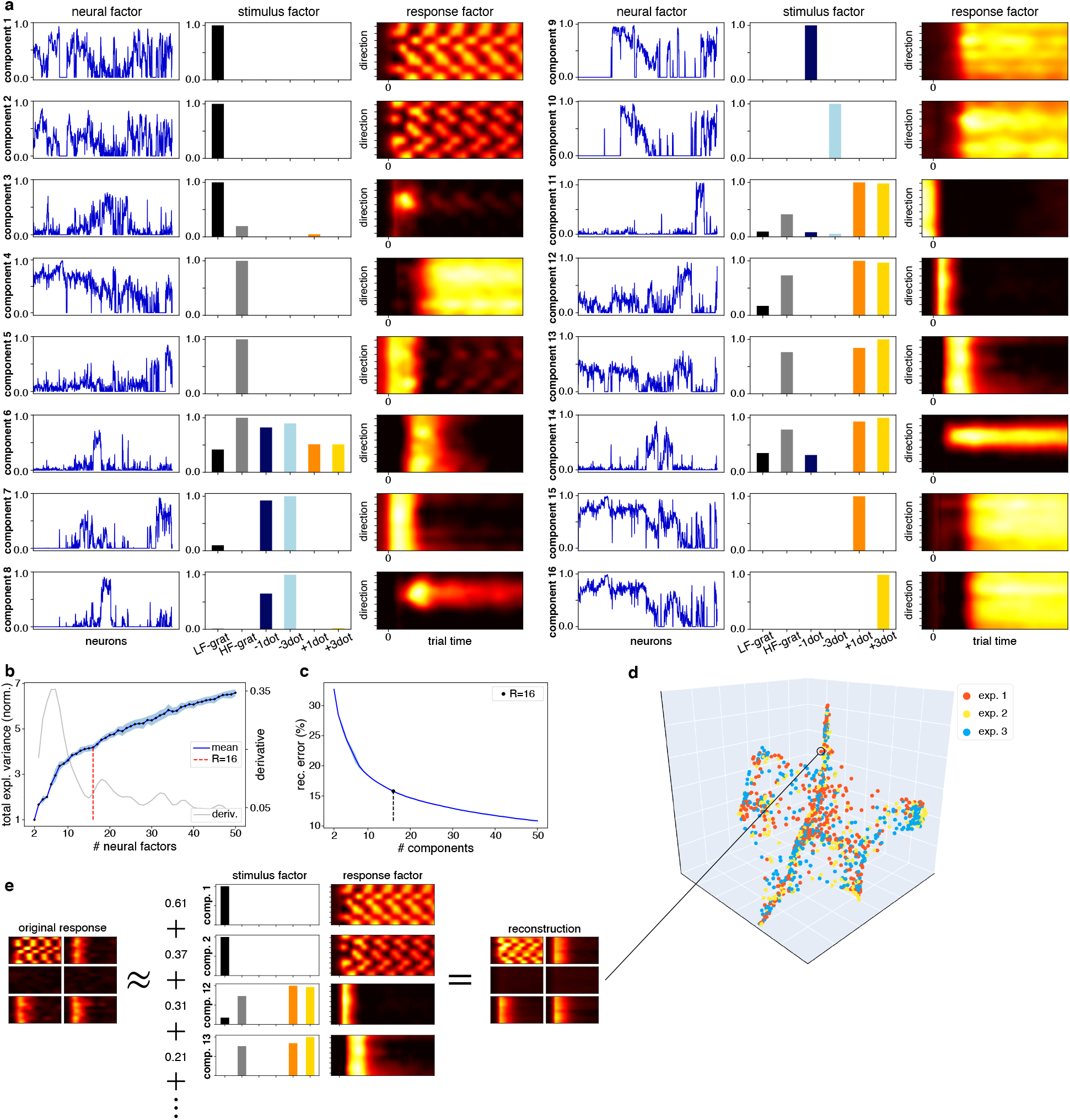
Tensor decomposition for the retinal data. **a**, The three factors comprising each component. Notice how the first three components are concentrated on low spatial frequency (LF) gratings, and how other components concentrate on high frequency (HF) gratings and the various flows. Among the activity factors, the first two indicate linear-like responses, while others are either transient, sustained, tuned, or delayed. Finally, notice how the neural factors are, in general, complex mixtures. **b**, Selection of appropriate number of components (*R* = 16) based on normalized explained variance (nuclear norm, see Methods, section 1.6). **c**, Reconstruction error as a function of the number of components (see Methods). **d**, Multiple recordings can be combined. Shown are data from three different experiments; neurons from different retinas are well-mixed across the manifold. **e**, Example of response reconstruction for a single cell using the components from **a**. Shown are the five components with largest neural factor loadings.

**Fig. S4:**
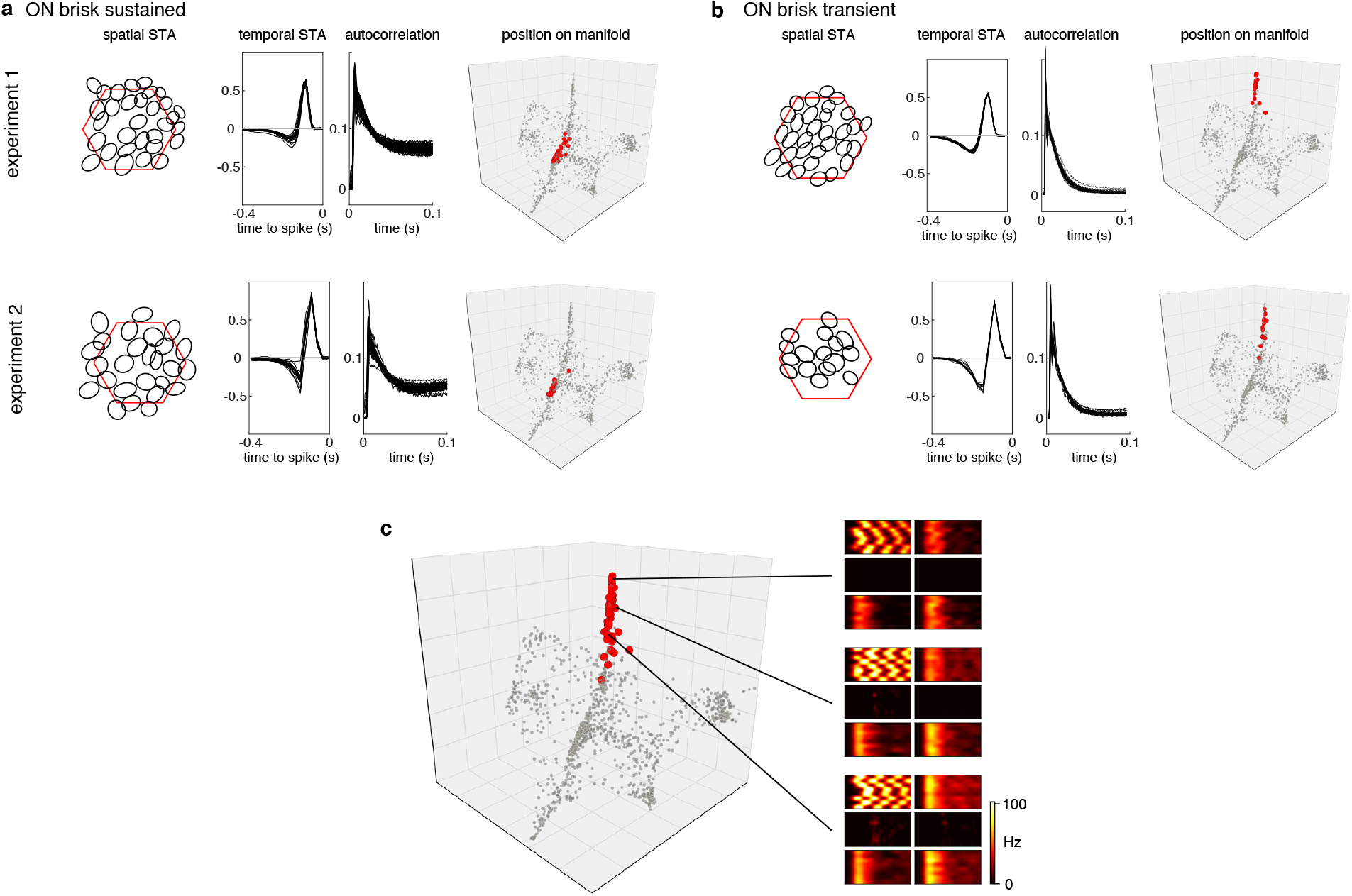
Understanding the retinal encoding manifold. **a–b**, Examples of retinal ganglion cells (RGCs) classified as ON brisk sustained (**a**) and as ON brisk transient (**b**) (data from two experiments). The classification is based on spatiotemporal spike-triggered averages (STAs) and spike autocorrelation curves, and was done independently (used a different stimulus) from building the encoding manifold (see Methods section1.1). Both of these RGC types are concentrated in specific regions (red dots) in the neural encoding manifold. **c**, ON brisk transient cells from all experiments combined in a single manifold (shown in red). Activity maps shown on the right reveal why these neurons cluster in an elongated fashion. Although qualitatively there is almost no change between them, there is a small, quantitative change in the relative magnitude of responses to high frequency gratings and positive flows compared to low frequency gratings. This slight increase, as one moves from top to bottom, is captured by the third (vertical) diffusion coordinate, highlighting an advantage of using a spectral embedding method (i.e., diffusion maps).

**Fig. S5:**
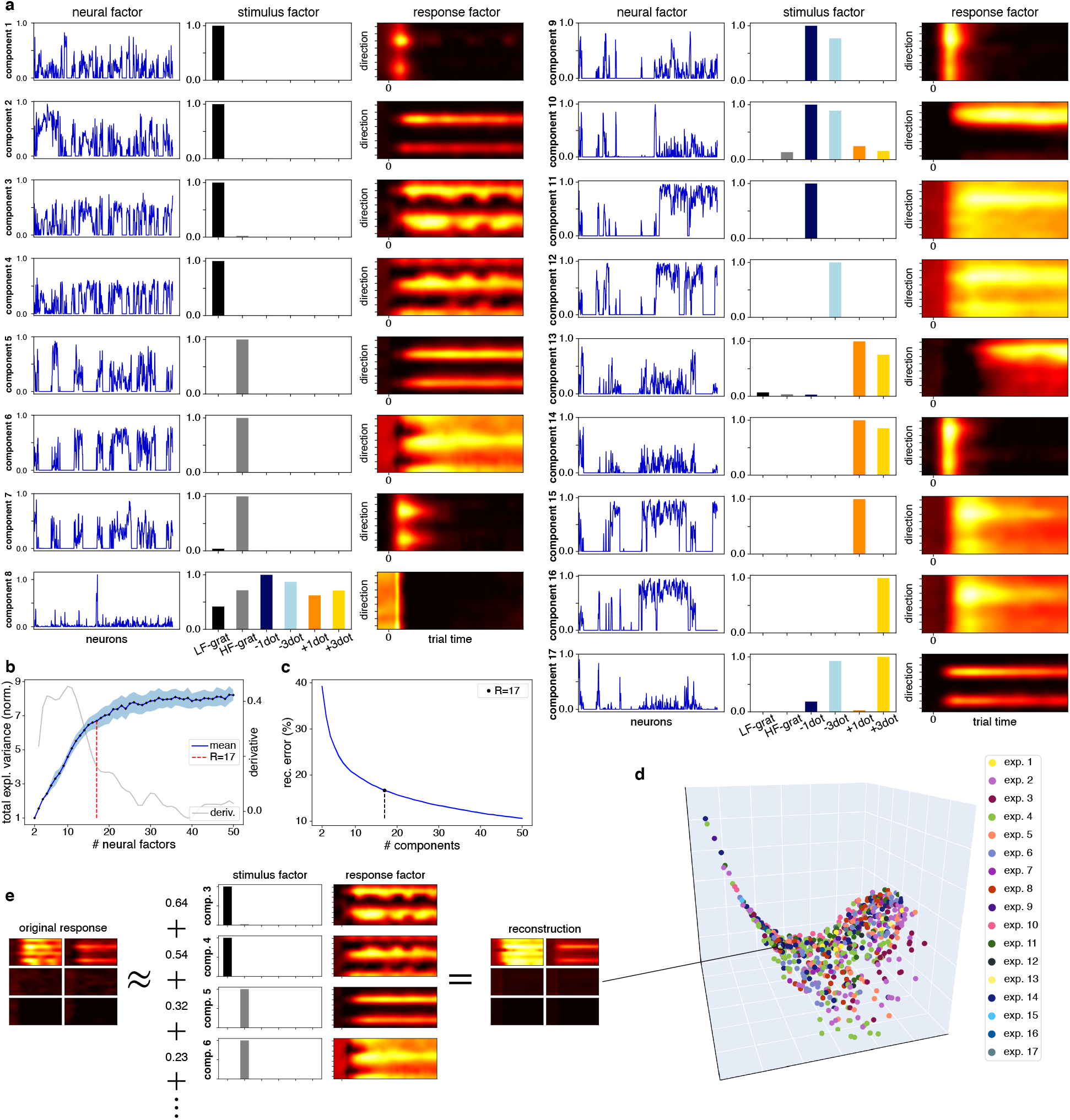
Tensor decomposition for the cortical data. **a**, The three factors comprising each component (compare with Fig. S3). Here we find response factors with various tuning widths, direction versus orientation tuning, as well as sustained versus transient temporal activity. **b**, Selection of appropriate number of components (*R* = 17, see Methods, section 1.6). **c**, Reconstruction error as a function of the number of components (see Methods). **d**, Different experiments are well-mixed across the manifold. **e**, Example of response reconstruction for a single cell as a weighted sum of the stimulus *×* response components from **a**, with weights given by the corresponding neural loadings (top four shown).

**Fig. S6:**
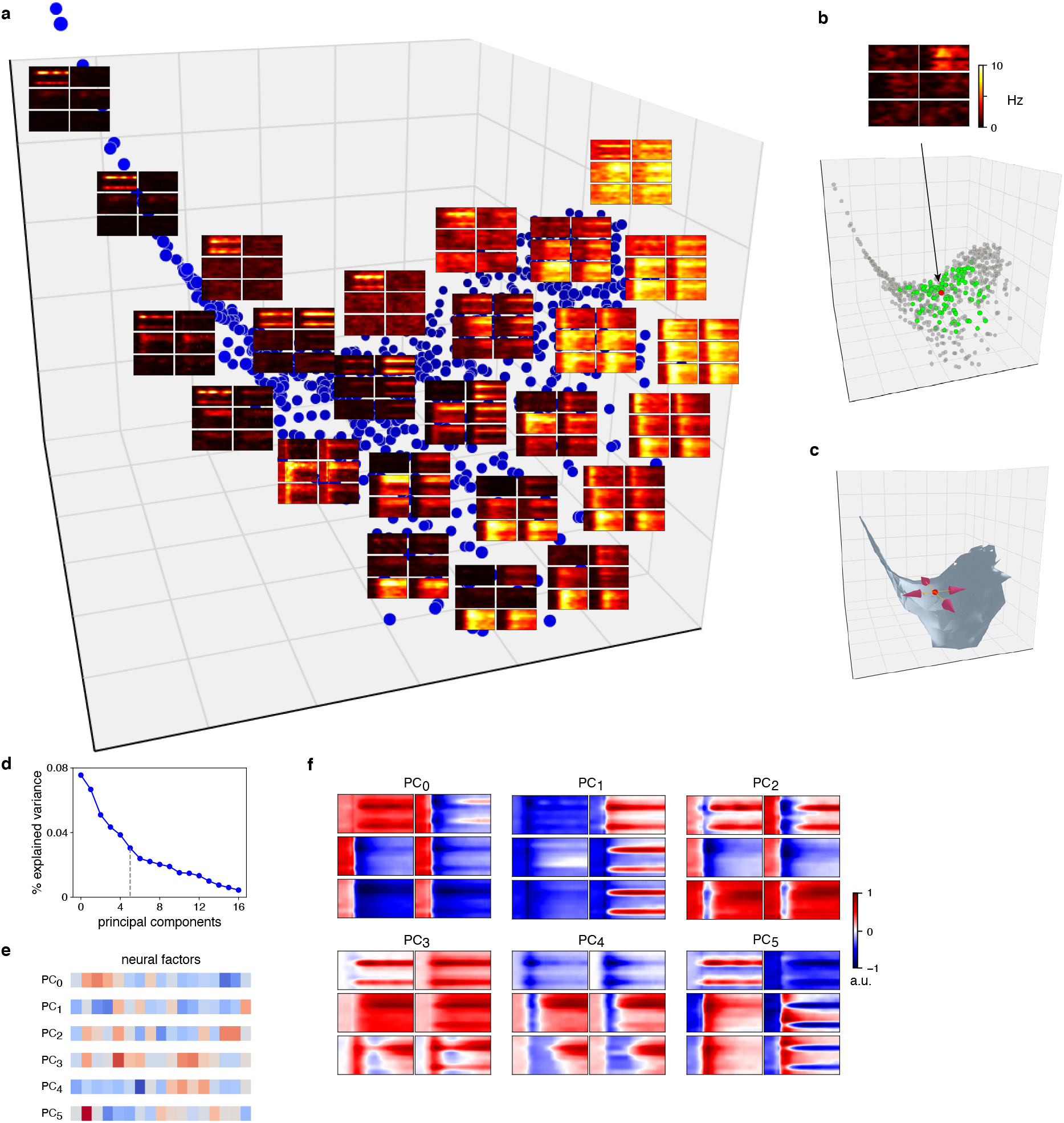
Understanding the cortical encoding manifold. **a**, Distribution of feature selectivity over the manifold. Here we show activity profile centroids (average response maps across a local neighborhood of 20 neurons) at 25 maximally distant positions (top view, as shown in inset) projected into the first three diffusion coordinates. Notice how selectivity (or visual feature preference) varies smoothly; there are no clear gaps as in the retinal manifold. The right corner shows neurons with little selectivity, and consists mostly of putative inhibitory neurons (cf. Fig. 4g). Cells responding exclusively to low-frequency gratings comprise the left arm, but responses to high-frequency gratings or flows gradually appear towards the center. The lower region is biased towards neurons responding strongly to positive contrast flows. Changing position across the manifold can be interpreted as how a neuron’s stimulus/response profile changes with respect to that of another neuron. These changes can occur in more than the three dimensions that are shown here, necessitating a dimensionality analysis. **b**, Illustration of dimensionality analysis (see Methods section 1.8). A neuron (*ν*) from the central region of the manifold, shown in red, has a local dimension of about 6 (Fig. 6), among the highest revealed by our stimuli. Nearby neurons specify a local neighborhood around *ν*, and have similar (but not identical) stimulus/response profiles. Mathematically, when the manifold is smooth, there is a tangent “plane” attached at *ν*, and moving across it indicates how the stimulus and response factors can vary as one moves a short distance along a dimension from *ν*. This view was discussed for the ring model in Methods, section 1.8, and is illustrated for V1 in **c**, showing arrows attached at *ν* over a continuous depiction of the cortical manifold. **d**, Performing local principal components analysis (PCA) on the neural matrix (see Methods, section 1.6) allows us to estimate these dimensions using *ν*’s local neighborhood in the data graph (neurons in green). Note how the spectrum agrees with the intrinsic dimension, and **e** how this procedure allows us to express each principal component (PC), or direction, in the neural encoding space. **f**, Because each neural factor is associated with a stimulus and a response factor, each PC can be visualized as a weighted sum of those. PC0 indicates a competition (positive versus negative values) between low-frequency spatial gratings and all other stimuli, and PC1 indicates a direction in which the orientation tuning for high frequency gratings is sharpened, in concert with oriented (3-dot) flows. PC5 indicates a preference for low frequency spatial grating in concert with single dot flows, but in competition with 3-dot flows.

**Fig. S7:**
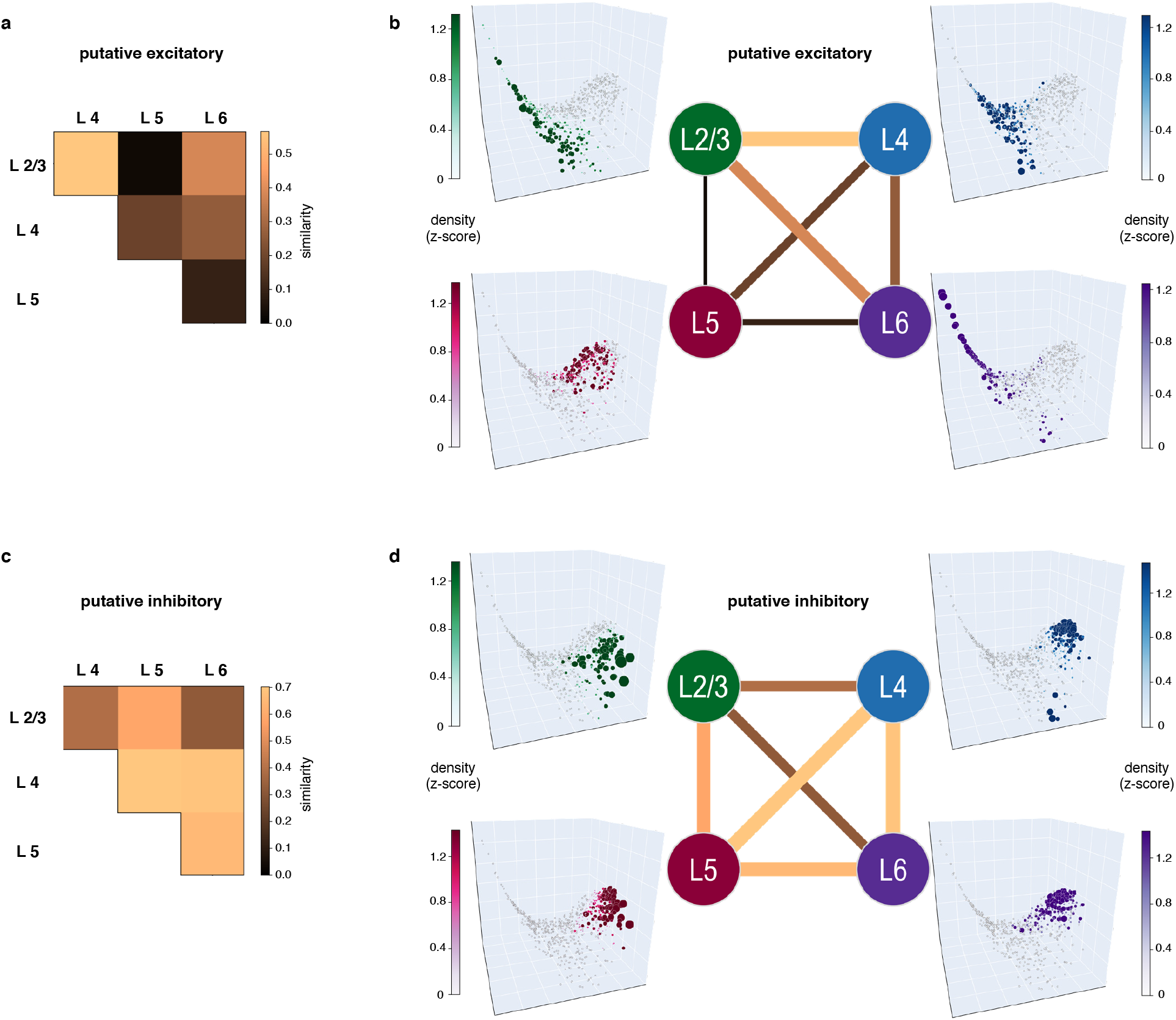
Physiological implications of the V1 manifold across layers. Encoding similarity across layers in V1, computed separately for putative excitatory (**a–b**) and putative inhibitory neurons (**c–d**) (see Methods, section 1.7). **a,c**, Similarities visualized as a matrix. Layers 2/3 and 4 were most similar for excitatory cells, in agreement with anatomical data from [42]. For inhibitory cells, layers 4, 5, and 6 were the most similar. **(b,d)** Similarities visualized as a graph in which layers are nodes and edges have thickness and color proportional to similarity, and corresponding layer densities (z-scores, see Methods) over the manifold. Because position on the manifold can be mapped back to response profiles (cf. Figs. 4 and S6), similarity between layers implies correlation between their overall feature selectivity, as indicated by responses to the stimulus ensemble. For example, for putative excitatory neurons (**b**), the high similarity between layers 2/3 and 4 follows from the common presence of neurons in both layers with narrow selectivity for gratings and/or flows (cf. Fig. S6), whereas layer 5 was the most distinct, comprised of mostly broadly selective cells (Figs. 4a,e and S6a), in agreement with data reviewed in [65]. For putative inhibitory neurons (**d**), we found roughly the opposite: feature selectivity in layer 5 was highly similar to the that found in other layers, while layer 2/3 had the most distinct selectivity due to the presence of many cells responding more strongly to the flow stimuli (towards the bottom of the manifold, cf. Figs. 4a,b and S6a).

**Fig. S8:**
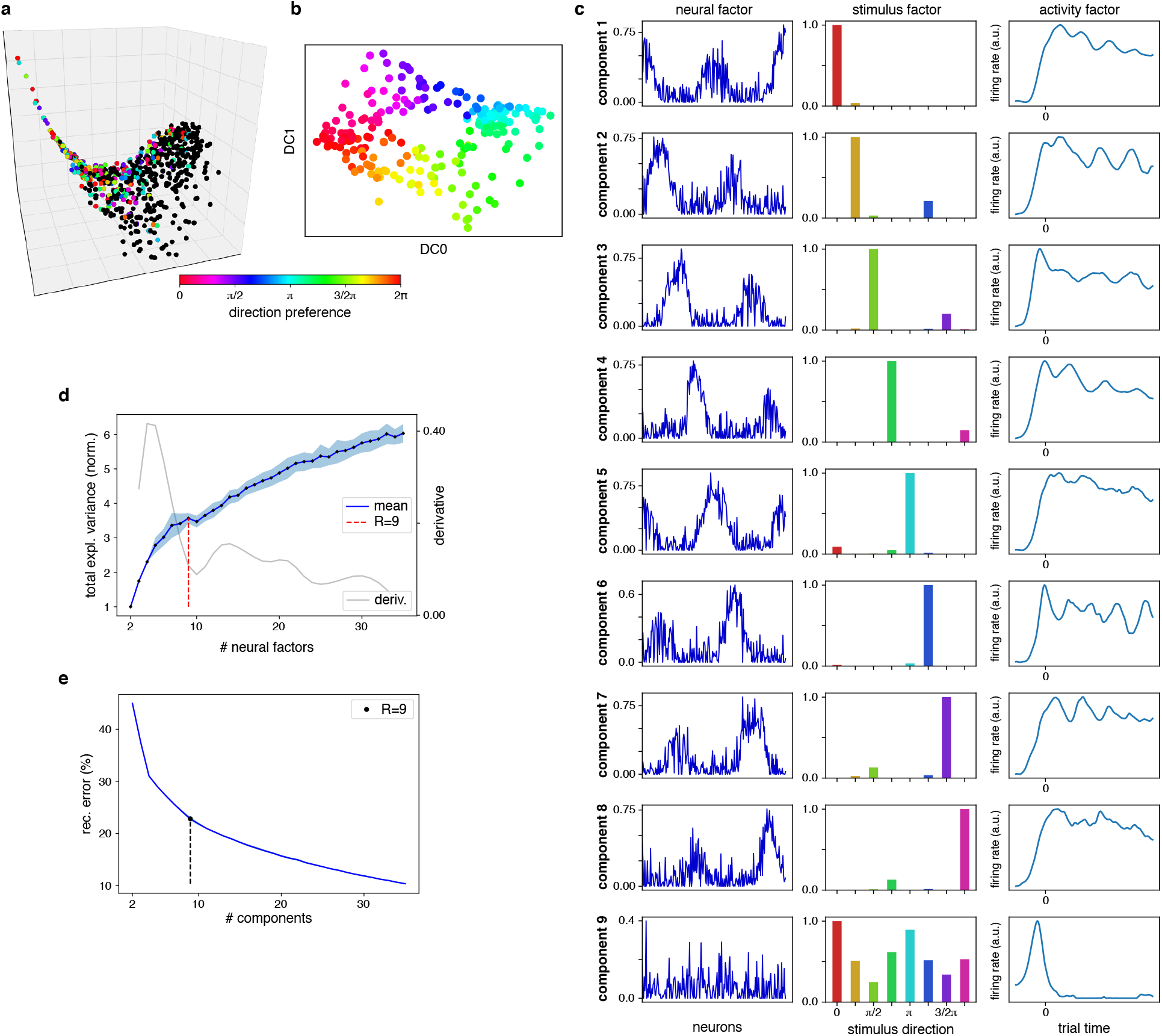
Organizing cortical responses to low spatial frequency gratings. **a**, We here illustrate the manifold inference process without the direction of motion normalization (see Methods). The dataset is restricted to cortical cells with well-tuned responses to the orientation or direction of low-frequency gratings (*n* = 225, labeled with colors corresponding to their preferred stimulus direction), and a tensor is built where the stimulus mode is comprised of the 8 directions of motion for low frequency gratings. **b**, The result is a ring-like manifold, in analogy with Fig. S1. **c**, By our algorithm, 9 components are produced. These reveal (in contrast with those from Fig. S1c) that most of the cells have periodic response curves. **d**, Although 8 stimulus directions were used, tensor decomposition requires a 9^th^ component (in contrast with the 8 for the artificial ring model, Fig. S2c) to account for variations in the latency period before the onset of activity. **e**, Reconstruction error as a function of the number of components (see Methods).

**Fig. S9:**
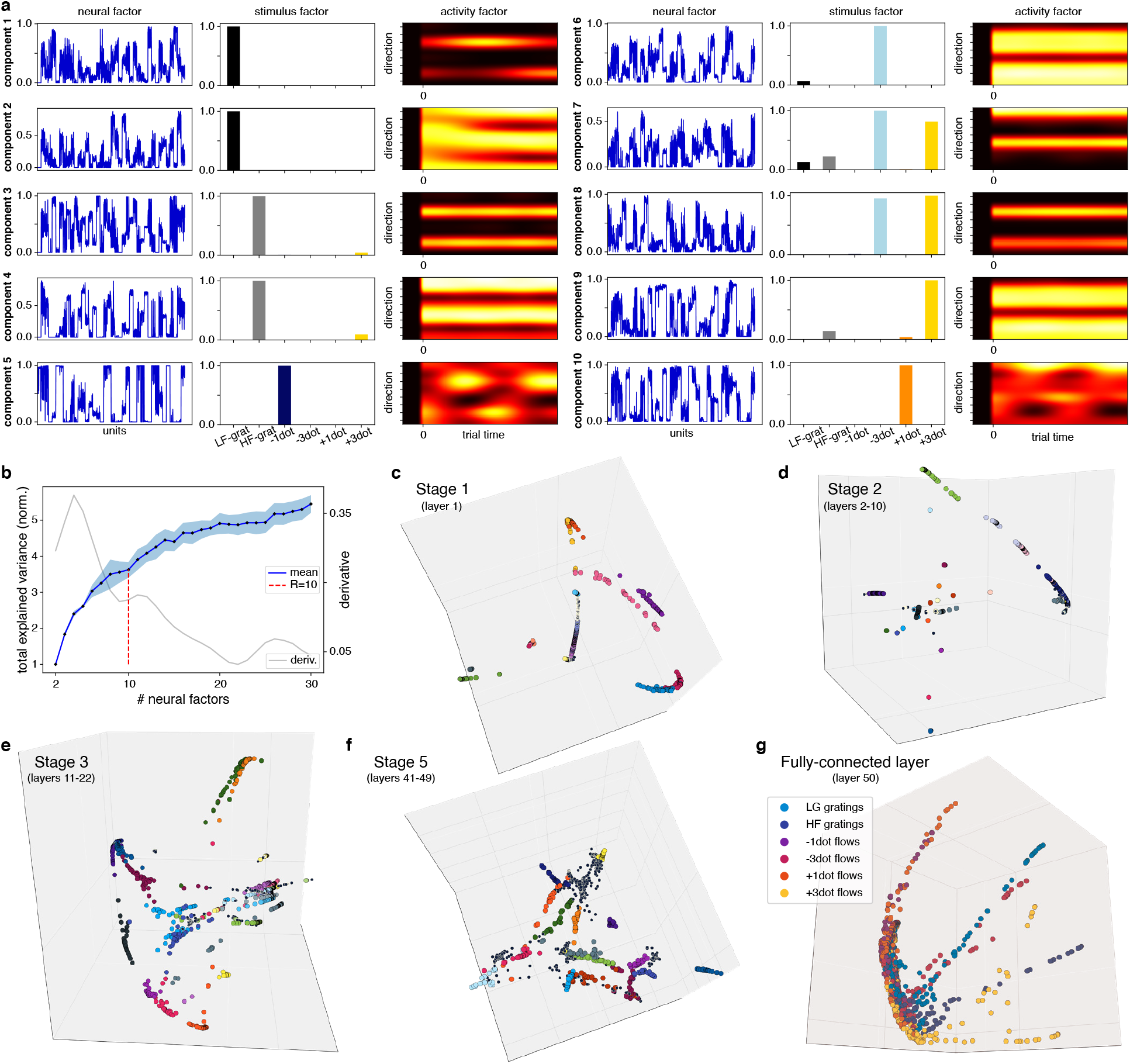
Tensor decomposition for ResNet50. **a**, The three factors comprising each component (compare with Figs. S3 and S5). Notice how the stimuli organize into separate factors, and how the neural factors reflect the block structure of the network. **b**, Tensor decomposition requires only 10 factors. **c–f**, Neural encoding manifolds for the remaining convolutional stages of ResNet50 (compare with Stage 4 in Fig. 5). Color labels represent the 40 feature maps with highest activation in response to our stimuli; neuronal units were sampled at random with probability proportional to their activation (see Methods, section 1.11). The topology remains largely disconnected throughout the network, even when individual layers are embedded separately (as in **c**). **g**, The fully-connected layer. Because there are no feature maps, colors now represent the stimulus with highest activation for each unit. Note the specificity of stimulus preferences, which corresponds to the classification performance.

**Fig. S10:**
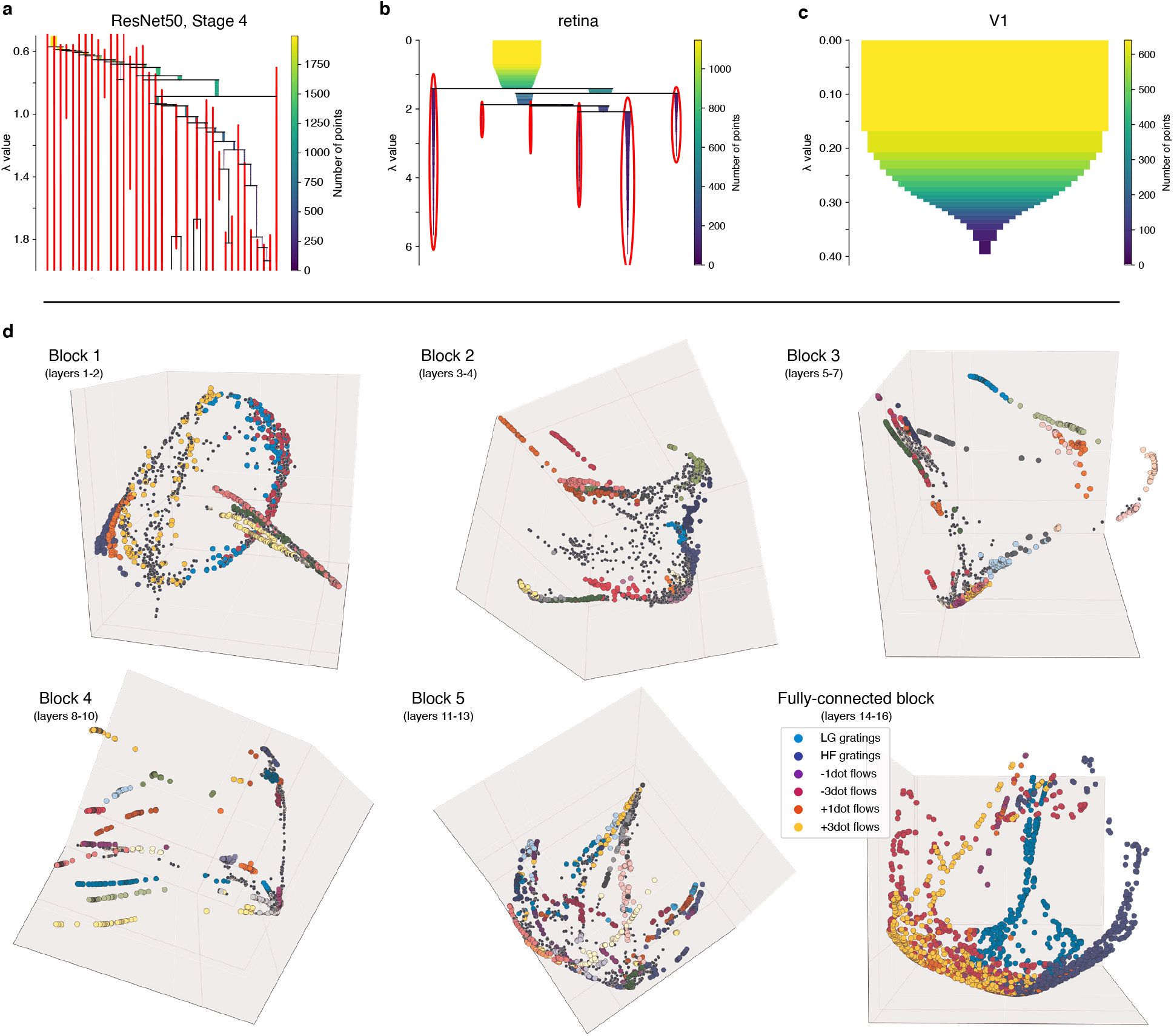
Hierarchical clustering (top) and encoding manifolds for VGG16 (bottom). **a**, Resulting condensed dendrograms from a density-based hierarchical clustering algorithm (see Methods, section 1.8) applied to ResNet50, stage 4 (cf. Fig. 5). A total of 34 clusters were identified, matching approximately the number of sampled feature maps. **b**, The clustered nature of the retina manifold is confirmed by the 6 identified clusters, while **c**, the continuous structure of V1 agglomerates all neurons together. **d**, Encoding manifolds for the different blocks of VGG16. The same descriptions made for ResNet50 (Fig. S9c–g) apply here.

